# Clean Enzymatic depolymerization of highly crystalline polyethylene terephthalate in moist-solid reaction mixtures

**DOI:** 10.1101/2020.07.06.189720

**Authors:** Sandra Kaabel, J. P. Daniel Therien, Catherine E. Deschênes, Dustin Duncan, Tomislav Friščić, Karine Auclair

## Abstract

Less than 9% of the plastic produced is recycled after use, contributing to the global plastic pollution problem. While polyethylene terephthalate (PET) is one of the most common plastics, its thermomechanical recycling generates a material of lesser quality. Enzymes are highly selective, renewable catalysts active at mild temperatures; however, the current consensus is that they lack activity towards the more crystalline forms of PET. We report here that when used in moist-solid reaction mixtures instead of the typical dilute aqueous solutions, enzymes can directly depolymerize high crystallinity PET in 13-fold higher space-time yield and a 15-fold higher enzyme efficiency than prior reports. Further, this process shows a 26-fold selectivity for terephthalic acid over other hydrolysis products, which allows the direct synthesis of UiO-66 metal-organic framework.

## Introduction

The amount of plastic manufactured annually is ever increasing, with 359 million tons produced in 2019.(*1*) Unfortunately, less than 9% of all plastic waste is currently recycled, owing to the lack of economically feasible, efficient and sustainable recycling technologies.(*2*) With an estimated 70 million tons produced annually,(*3*) polyethylene terephthalate (PET) is one of the most common consumer plastics, making up disposable bottles and polyester clothing fibers, among others. Current recycling of PET, typically achieved through thermomechanical means, significantly reduces plastic integrity,(*4*) thereby limiting the applications of recycled PET to lesser quality end-of-life products that are eventually discarded (*e*.*g*. carpets).(*5*) Consequently, better PET recycling technologies are urgently needed to move towards a circular economy that minimizes the accumulation of plastic in landfills and the environment.(*2*)

One alternative to thermomechanical recycling involves depolymerizing PET into its building blocks, terephthalic acid (TPA) and ethylene glycol (EG), which can then be recycled to produce high-quality PET products by re-polymerization (*i*.*e*. true recycling).(*6*) Chemically breaking down PET typically involves the use of a strong acid (*e*.*g*. sulfuric, nitric, or phosphoric acid) or base (*e*.*g*. NaOH or KOH), in combination with elevated temperatures and/or pressures, and with the production of large volumes of toxic waste.(*7–11*) Biocatalysts capable of hydrolyzing PET offer a more sustainable route to plastic recycling, as these environmentally benign, renewable catalysts can work under mild conditions and generate a clean product without the use of hazardous chemicals. Several enzymes have been reported that are capable of depolymerizing low crystallinity PET (<20%), including the remarkably efficient mutant of leaf-branch compost cutinase recently developed by Tournier *et al*.;(*12*) however, the efficient depolymerization of highly crystalline forms (>20%) of PET typically found in consumer products (Table S9) has not been reported.

Herein we show that switching from standard dilute aqueous environments to unconventional moist-solid reaction mixtures enables the direct enzymatic depolymerization of high crystallinity PET, thus avoiding the otherwise necessary thermal or chemical pre-treatment. When applied to the thermostable *Humicola insolens* cutinase (HiC), marketed as Novozym® 51032, this unique approach yields TPA selectively, with both crystalline and amorphous PET regions hydrolyzed to the same extent. The purity of TPA is sufficiently high to permit conversion to value-added products, such as the microporous metal-organic framework (MOF) UiO-66. Overall, we demonstrate that enzymatic PET depolymerization in the absence of solvent is not only more efficient, but also proceeds with higher selectivity and with considerably improved space-time yield (STY) and enzyme efficiency compared to prior reports (Table S9).

## Results

### Enzymatic hydrolysis of PET by HiC in moist solids

Commercial PET powder of 36% crystallinity, similar to a post-consumer PET bottle, was used as a model substrate to develop our process (Figs. S2, S3). The reactions were typically carried out using a mechanoenzymatic approach,(*13*) based on an initial brief period of ball milling (5 min milling at a frequency of 30 Hz) (*14–16*) in the presence of enzyme (0.6 wt%) and a minimal amount of buffer (0.1 M sodium phosphate, pH 7.3, at a liquid-to-solid ratio *η* = 1.5 μL mg^−1^), followed by 7 days of aging (static incubation) at 55°C (*17–19*). This process gives reaction mixtures with paste-like consistency, which progress to a yield of 20 ± 1% TPA (Fig. 1A–D, Table S1, entry 4). HPLC analysis of the crude mixture revealed a 20-fold selectivity for TPA over mono(2-hydroxyethyl) terephthalate (MHET) and no detectable amount of bis(2-hydroxyethyl) terephthalate (BHET) or other oligomers (Fig. 1D). Remarkably, manual mixing of the reactants instead of ball milling was enough to obtain TPA in 19 ± 1% yield within 7 days (Fig. 2A, Table S1 entries 1–6). To ensure reproducible mixing of the reactants, however, we continued to use ball milling in subsequent experiments. Attempt to conduct the reaction under identical conditions, but in the absence of the enzyme, led to only 0.02% conversion to TPA (Table S1 entry 25). As ball milling was observed to increase the crystallinity of PET (*vide infra*), these results reveal that no significant amorphization or hydrolysis of PET takes place due to milling or aging alone. Lastly, to eliminate the possibility that other components of the commercial enzyme solution (*e*.*g*. cryoprotectants, salts) participate in the reaction, a buffer-exchanged lyophilized enzyme powder was prepared and tested for activity. This cleaner enzyme preparation led to a comparable 18 ± 3% yield of TPA (Table S2 entry 1).

**Fig. 1.**
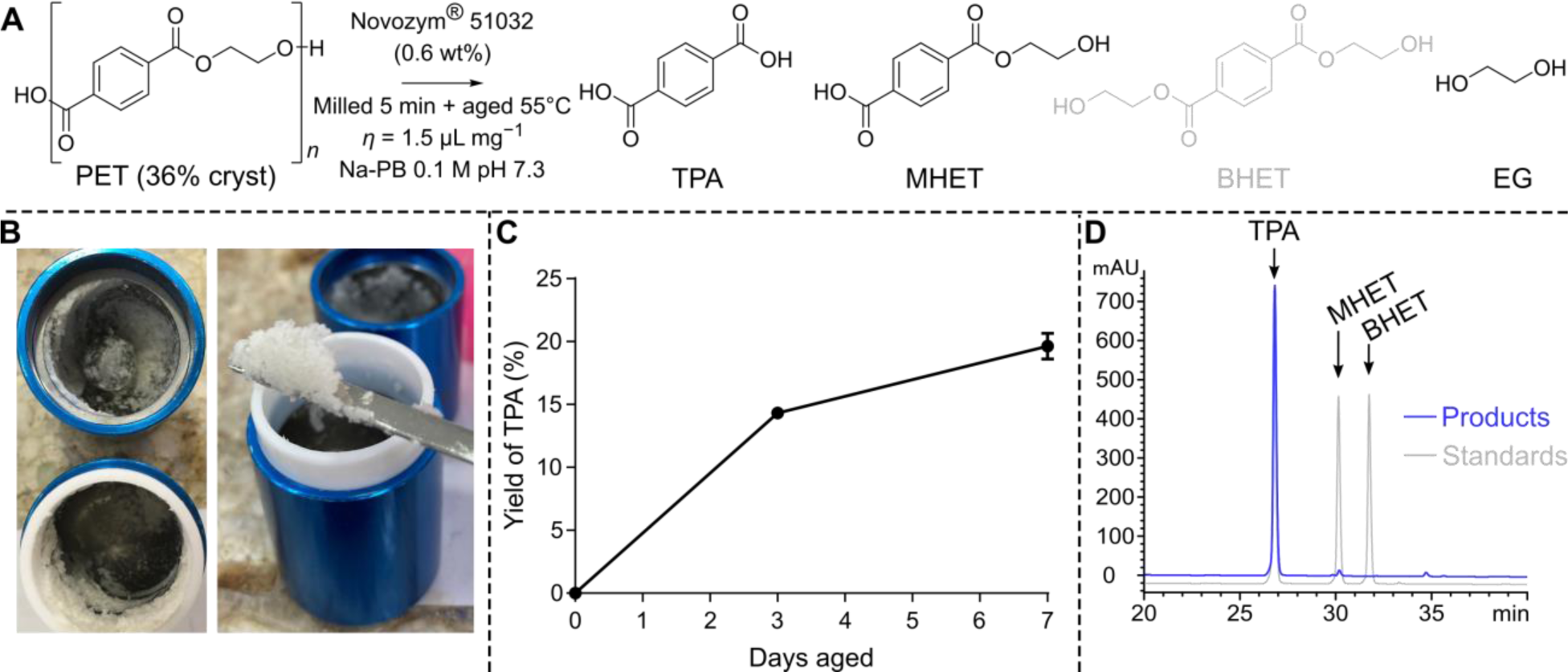
A) General reaction scheme of the mechanoenzymatic hydrolysis of PET using HiC (Novozym® 51032), showing the possible hydrolysis products terephthalic acid (TPA), mono(2-hydroxyethyl) terephthalate (MHET), bis(2-hydroxyethyl) terephthalate (BHET) and ethylene glycol (EG). BHET (in gray) was not detected. B) Photos of the paste-like reaction mixture after 5 minutes of milling. C) The yield of TPA (% of theoretical) achieved during static aging at 55°C, after 5 minutes of initial ball milling. D) Chromatograms of the reaction products (blue) compared to traces of standard samples of TPA, MHET and BHET (gray) monitored at 240 nm.

**Fig. 2.**
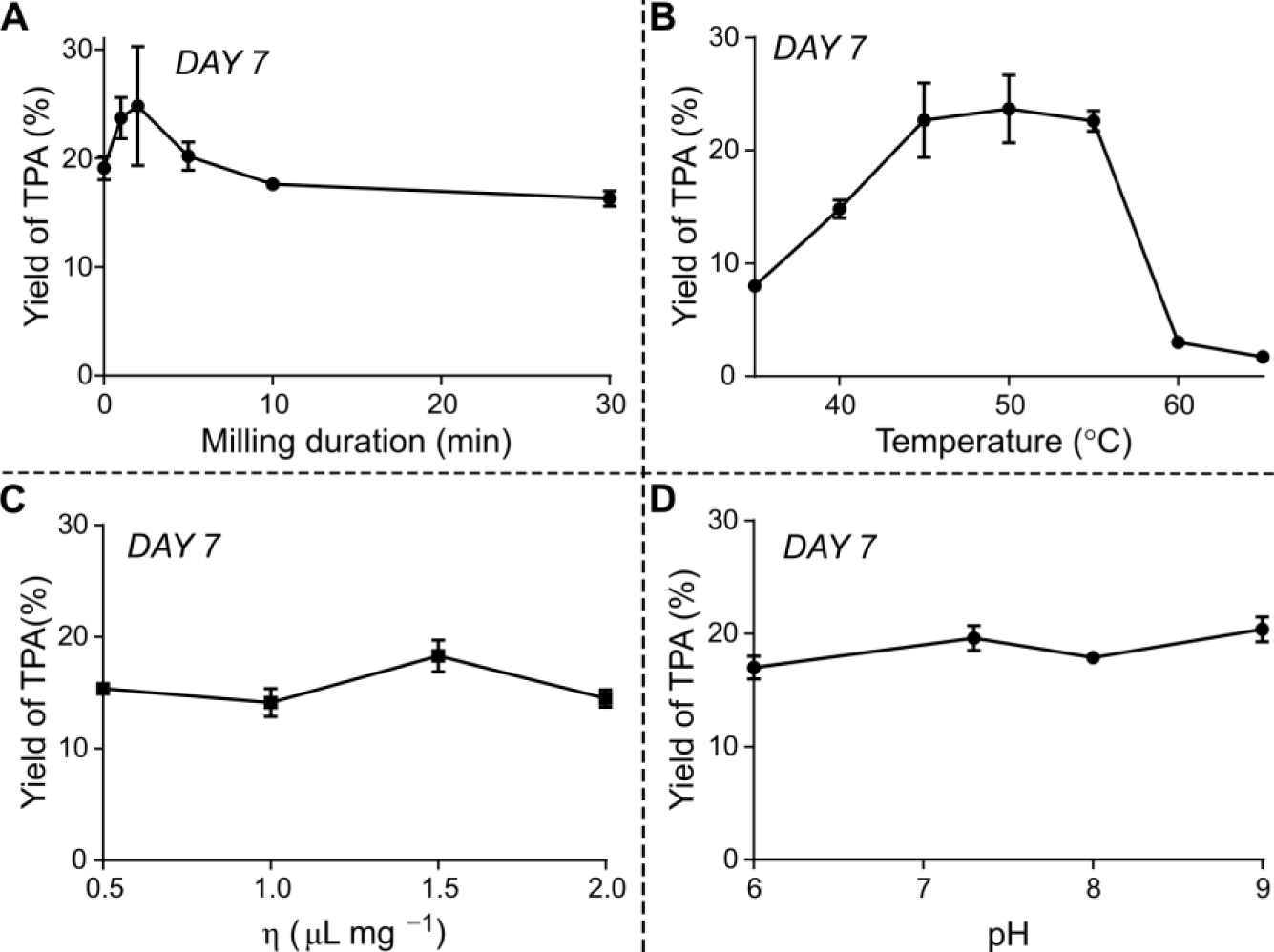
Reaction outcome (day 7) depending on the duration of the initial milling where 0 min is manual mixing (A), at different aging temperatures (B), at different liquid-to-solid ratios (C) and at different buffer pH (D).

The optimal aging temperature was found to be 45–55°C (Fig. 2B, Table S1 entry 7–13), a temperature range easily accessible, for example, in a greenhouse. Notably, in-solution depolymerization of PET by HiC is efficient only at temperatures above 60°C, and only with low crystallinity PET (Table S9). All aging experiments were performed in closed vessels to avoid drying of the reaction mixtures. The steep decline of activity at aging temperatures ≥ 60°C may indicate that the thermal stability of the enzyme in moist-solid reaction mixtures is diminished, as HiC activity in solution was reported to persist even at 70°C (Table S9).(*20*) Varying the liquid-to-solid ratio between 0.5 – 2 μL mg^−1^ demonstrated that HiC may function well with even less water than under our preferred conditions (Fig. 2C, Table S2 entries 6–9). Using less water affords similar yields, but with an improved STY after 7 days at 55°C (1.58 g_TPA_ L−1 h−1). This is remarkable, considering that a liquid-to-solid ratio *η* = 0.5 μL mg^−1^ corresponds to only 2-fold molar excess of water in the HiC-catalyzed PET hydrolysis.

Based on its poor water solubility (0.015 mg mL^−1^ at 20°C), we did not expect TPA to dissolve significantly in the buffer. Consistent with this expectation, powder X-ray diffraction (PXRD) patterns recorded after 7 days of aging at 55°C (Fig. S4) show the presence of crystalline TPA within the solid reaction mixture. The pH of the buffer is not expected to be meaningfully affected by the presence of crystalline TPA. Indeed, the HiC enzyme maintained a similar level of activity over a pH range of 6–9 (Fig. 2D, Table S1 entries 4, 15–17), regardless of the nature and concentration (0.1 – 1 M) of the buffer (Table S1 entries 19, 20). Reactions carried out with water only, however, gave a somewhat reduced TPA yield of 14 ± 1% (Table S1 entry 18).

### HiC-catalyzed mechanoenzymatic hydrolysis of post-consumer PET and other plastics

To explore the efficacy of our process on PET from other sources, we first investigated the hydrolysis of a powdered, transparent post-consumer PET bottle (30% crystalline, see SI). Ball milling (30 minutes) and aging (7 days) of this PET powder with 0.6 wt% of HiC, with *η* = 1.5 μL mg^−1^ of 0.1 M sodium phosphate buffer (pH 7.3) gave a 16 ± 2% yield of TPA (Table S3 entry 1). Notably, a comparable yield of TPA (15 ± 4%) was also achievable by milling the intact PET bottle pieces with the enzyme for a longer time (30 minutes) before aging, suggesting that pre-micronization of the sample could be avoided altogether (Table S3 entry 2).

An equal yield of TPA was achieved when the bottle PET was pre-milled with 10 wt% of polypropylene cap pieces (Table S3 entry 3). Similarly, a powdered black post-consumer PET container, labelled as containing 80% recycled content, yielded 15 ± 1% TPA under similar conditions (Table S3 entry 4). This result further establishes that PET hydrolysis in moist-solid environment can proceed even in the presence of common post-consumer plastics impurities. Non-participating polymers have previously been reported to increase the activity of *β*-glucosidase under mechanochemical conditions.(*21*) Testing the effect of other polymer additives on PET depolymerization by HiC yielded similar TPA yields (16% with microcrystalline cellulose and 19% with polystyrene), suggesting that polymeric impurities may not affect enzyme activity (Table S4).

To our surprise, pre-milling of amorphous thin film PET (13% crystallinity) increased crystallinity to 33%, as established by differential scanning calorimetry (DSC, see SI). This observation was attributed to stress-induced crystallization,(*22*) which prevented us from obtaining low-crystallinity PET in powdered form. A reaction on the pre-milled powdered thin film resulted in 17 ± 1% yield of TPA (Table S3 entry 5), similar to what is observed with other PET materials of similar crystallinity.

The reactivity of HiC was also explored on polyester textiles, which were efficiently powdered during the 5 minutes milling step at the start of the reaction. For example, a dark blue textile labelled as 100% polyester afforded 6.3 ± 0.8% of TPA after 5 minutes of milling and 7 days of aging (Table S3 entry 6). We suspect that other undisclosed constituents or textile finishing treatments may lead to reduced or underestimated yields.

Lastly, we explored the scope of our process for the HiC-catalyzed depolymerization of other plastics, including polybutylene terephthalate (PBT), polycarbonate (PC), and polycaprolactone (PCL). Gratifyingly, even with mechanoenzymatic reaction conditions optimized for PET, modest depolymerization of these plastics was achieved, consistently outperforming comparable in-solution reactions (Table S5).

### Acceleration of PET hydrolysis by reactive aging (RAging)

Previous reports on the mechanoenzymatic breakdown of the biopolymers cellulose,(*21,23, 24*) xylan(*25*), and chitin(*26*) demonstrated that enzyme kinetics under aging conditions are hyperbolic. These reactions can be accelerated by periodically repeating the milling process just before the yield reaches a plateau, in a process termed reactive aging (RAging).(*23*) The RAging cycles were designed to take advantage of the initial faster reaction rates of aging and improved mixing due to repeated milling.

Kinetic analysis of the HiC-catalyzed PET depolymerization (after 5 minutes of milling) revealed an initial rate of 10.4 mM_TPA_ h−1, that decreased rapidly after 24 h (Fig. 3A). Given this information, we elected to use RAging cycles consisting of 5 minutes ball milling at 30 Hz and 24 h aging at 55°C. This reaction design led to a 21 ± 2% yield of TPA in only 3 days (Fig. 3B, Table S6 entry 1), compared to 7 days needed to achieve the same yield if milling was performed only once. The RAging reaction was also successfully scaled up 3-fold, using the same 15 mL milling jars, without any decrease in conversion (Table S6 entry 2). While RAging was clearly advantageous to speed up the depolymerization, the reactions typically converged at 20–25% yield (Fig. 3B).

**Fig. 3.**
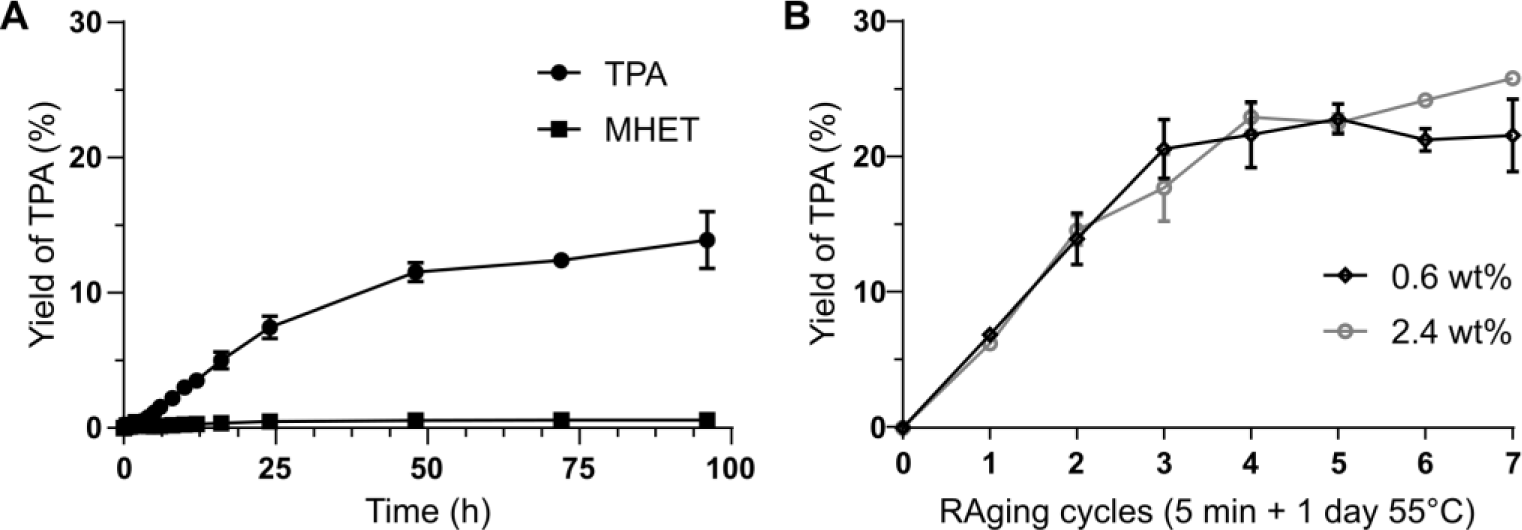
(A) The kinetics of HiC (Novozym® 51032) during aging at 55°C for the mechanoenzymatic hydrolysis of PET powder (36% crystallinity). (B) Formation of TPA (%) during RAging reactions at 0.6 wt% (black) or 2.4 wt% (gray) HiC loading.

### Mechanistic considerations

Next, we systematically explored possible explanations for the yield to plateau at 20-25%. Increasing the enzyme loading 4-fold brought no improvement in TPA yield (Fig. 3B, Table S6 entry 3). Product inhibition was ruled out given that PET hydrolysis proceeded to the same extent even in the presence of 18 wt% of TPA and/or EG added from the start of the reaction (Table S7 entries 1–3). This result also confirms that the reaction convergence near 25% yield is not caused by TPA-induced pH changes, as the content of TPA in these reaction mixtures is doubled at the end of the process.

Notably, DSC analysis of the remaining PET after removal of the hydrolysis products showed that PET crystallinity did not differ significantly from the native material (Fig. S2), indicating that both crystalline and amorphous PET regions were hydrolyzed to the same extent. This is unexpected, and in stark contrast with enzyme activity in solution which preferentially depolymerize the amorphous PET regions.(*20, 27*–*32*) Similarly, PXRD (Fig. S4) and Fourier-transform infrared spectroscopy (FTIR, Fig. S5) analysis of the remaining PET showed no detectable changes to the chemical or surface structure of the material. Therefore, it is unlikely that the plateau near 25% reaction yield is due to depletion of amorphous PET and concentration of the more resistant crystalline material on particle surfaces. Scanning electron microscopy (SEM) images revealed that the smooth surface of native PET particles (Fig. 4A) became rougher after mechanoenzymatic hydrolysis (Fig. 4B), but without any significant pitting.

**Fig. 4.**
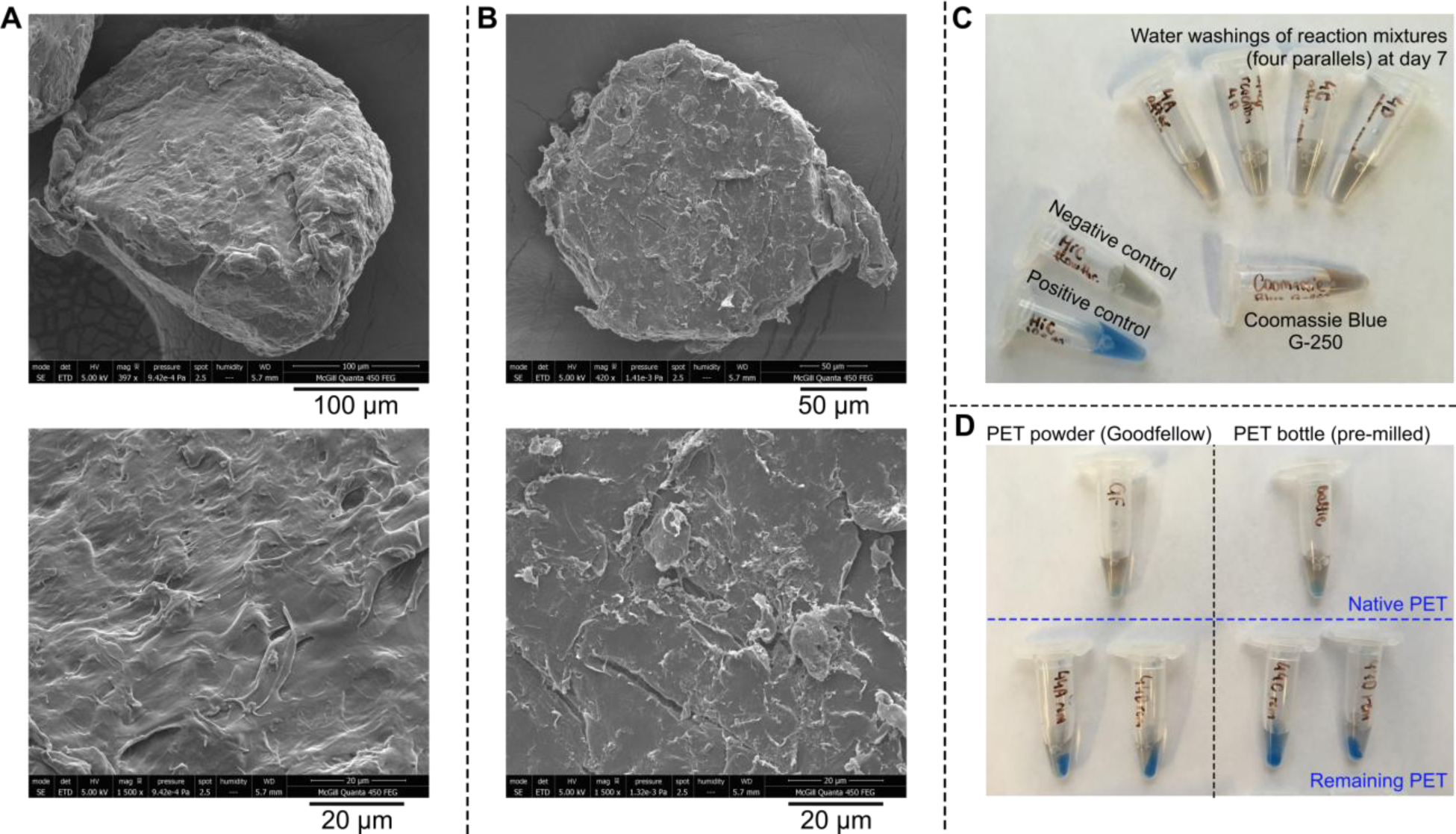
SEM micrographs of the PET particles before (A) and after (B) the HiC-catalyzed mechanoenzymatic reaction. Particles shown after the reaction were cleaned of the hydrolysis products by washing with methanol (see SI). Panels C and D show the results of protein detection using the Coomassie Blue G-250 dye. Panel C shows that none of the HiC can be extracted by water at day 7, as the washings did not stain blue. The positive control was obtained from the commercial enzyme solution at a concentration equal to that potentially extracted from the reaction mixture, and the negative control was performed on the flow-through from the commercial enzyme solution after using a 10 KDa MWCO centrifugal concentrator to selectively remove the enzyme. The color of unreacted Coomassie Blue is also shown for comparison. Panel D shows qualitative staining result for native PET alone (above) and the washed remaining PET after HiC-catalyzed mechanoenzymatic reactions (below). Remaining PET from two washing conditions (Method A and B, see SI) are shown for both PET powder (left) and pre-milled bottle PET (right).

We next attempted to extract the enzyme from the paste-like reaction mixture at various time-points to measure its remaining activity. HiC was easily extracted into buffer after 5 minutes of milling and the extract exhibited full activity, showing that our milling conditions are not detrimental to the enzyme. In contrast, no active enzyme could be extracted from the samples after 3 or 7 days of aging (see Table S8). In fact, based on a Coomassie Blue staining test, no detectable protein moved to the buffer from the solid paste after aging (Fig. 4C). Instead, even after subsequent separation of the hydrolysis products (see SI section *TPA isolation*), most of the protein remained on the PET surface (Fig. 4D). Subjecting this solid residue to a BHET enzyme activity assay revealed that the enzyme present on the PET surface exhibits only about 4% of its original activity (Table S8 entries 8 and 9). This dataset allows us to conclude that after 3–4 days of reaction, most of the protein is denatured and adsorbed on PET.

We therefore sought to establish the cause of HiC denaturation. Keeping HiC in buffer at 55°C for 3 days prior to the mechanoenzymatic reaction did not reduce enzymatic activity (Table S1 entries 22 and 23), implying that the enzyme denaturation observed during PET hydrolysis is not induced by temperature. To explore the relationship between enzyme activity and binding to hydrophobic polymer surfaces, the activity of HiC was evaluated after aging with polybutylene terephthalate (PBT), a material that structurally resembles PET but is a poorer substrate. Only ca. 1% PBT hydrolysis was observed after 3 or 7 days of aging under the same conditions employed for PET, and HiC extracted from this reaction mixture retained 65% activity (BHET assay, Table S8). These results indicate that the active enzyme does not bind tightly to the surface of hydrophobic polymers and confirms that strong attachment to PET is specific to the denatured HiC enzyme. Furthermore, as active HiC is readily extracted from the surface of the poor substrate PBT, but not from the surface of the much more reactive PET, we conclude that the observed plateau in reaction yield results from activity-induced denaturation of the enzyme – which may be overcome by switching to a different enzyme or by protein engineering.

### Maximizing the yield of TPA from PET

Given the above results, it was envisaged that the TPA yield might be improved by replenishing the denatured enzyme once the reaction reaches a plateau (while keeping the liquid-to-solid ratio constant at *η* = 1.5 μL mg^−1^). Commercial HiC is available as a solution, which means that each enzyme addition increases *η* by 0.3 μL mg^−1^, which may be detrimental to the reaction (Fig 2C). For this reason, the PET remaining after the first 3-cycle RAging experiment (which had reached a TPA yield of 17.0 ± 0.6%) was washed and dried before being submitted to another 3-cycle RAging process with fresh enzyme and buffer added (enzyme loading 0.6 wt% towards the remaining PET). This overall process afforded an additional 10 ± 1% yield. Repeating rounds of 3-cycle RAging experiments that include PET washing, followed by addition of fresh enzyme and buffer, all proceeded with an approximate 10% additional yield of TPA from the remaining PET (Fig. 5A), allowing us to reach an overall 49 ± 2% yield of TPA from 900 mg of PET over 7 rounds, using a total of only 3 wt% of enzyme (Fig. 5B).

**Fig. 5.**
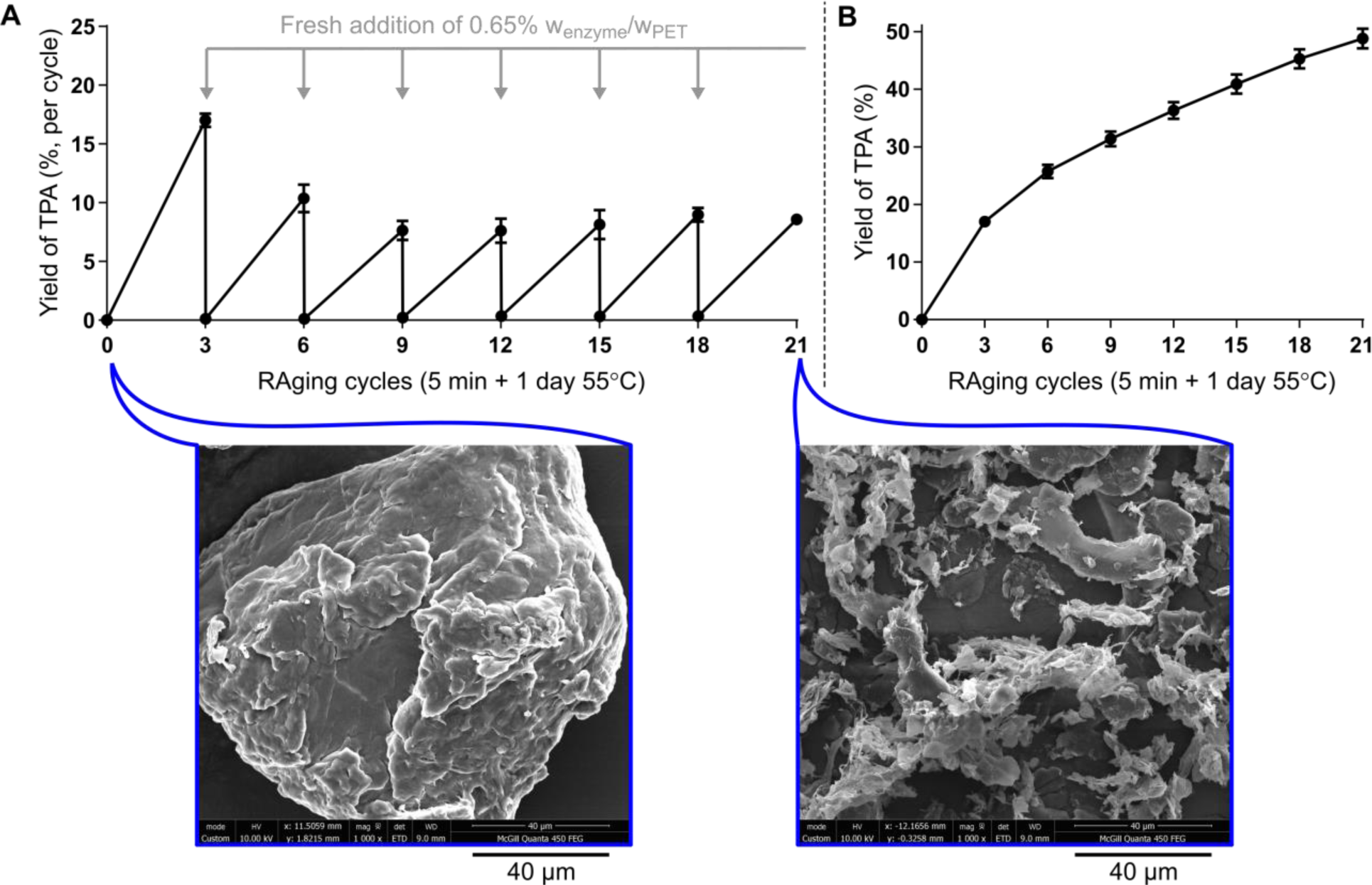
Per-round (A) and cumulative (B) yield of TPA in the multi-round RAging experiments. More enzyme (0.65 wt%) and enough buffer to ensure a constant liquid-to-solid ratio were added after every round of RAging (3 cycles), and the obtained TPA was extracted. The reaction continued with the remaining washed and dried PET. SEM micrographs are shown (bottom) for the isolated and washed PET substrate both at the start of the reaction and for the remaining PET after the multi-round experiment.

The reason for the drop in PET conversion past the first round is not clear and may be influenced by several parameters (enzyme adsorption, reaction kinetics, additives in the plastic, etc.). As mentioned above, our current understanding is that enzymatic catalysis itself hastens the denaturation of HiC, which then unfolds and sticks to the remaining PET substrate over time. Enzyme retention on the PET surface, after the removal of products and washing, was confirmed by energy dispersive X-ray spectroscopic analysis (EDS), showing the presence of nitrogen in the sample after the first RAging cycle (Fig. S9) and also after the last cycle (Fig. S10). In contrast, nitrogen was not present in non-hydrolyzed PET (Fig S8). The size of PET particles remaining after the entire RAging process is diminished, with their surface exhibiting increased roughness (Fig. 5 bottom).

### Isolation of the recovered TPA and its use in the synthesis of a metal-organic framework (MOF)

The TPA produced by mechanoenzymatic hydrolysis of PET is readily isolated from the reaction mixture by washing with methanol, or more conveniently with aqueous sodium carbonate to obtain a water-soluble salt from which TPA can be subsequently precipitated by acidification (see SI for further details). The sodium carbonate solution was used to isolate all soluble reaction products between rounds in the multi-cycle experiment (Fig. 5A). Analysis of the products by PXRD (Fig. S4), FTIR (Figs. S6, S7), and ^1^H NMR (Figs. S11–S13), revealed that the isolated TPA is not only crystalline but also >95% pure, with the only minor product detected being MHET (up to 4 wt%). Recently, Tournier et al. demonstrated that TPA obtained by enzymatic depolymerization of PET waste can be used to make like-new PET bottles with equal mechanical properties to those of the petrochemically derived PET.(*12*) Alternately, TPA derived from PET by chemical hydrolysis with strong bases or acids has been used to make value-added materials, such as MOFs.(*33*) By adapting a reported mechanochemical method,(*34, 35*) we converted the herein obtained TPA to the popular and commercially significant MOF UiO-66. The synthesis of UiO-66 was accomplished by ball milling (20 minutes, 30 Hz) TPA with the pre-synthesized dodecanuclear zirconium acetate cluster Zr_12_O_8_(OH)_8_(CH_3_COO)_24_ in the presence of a small amount of methanol (*η* = 0.66 μL mg−1) in a 15 mL ZrO_2_ milling jar containing a single 3.2 g ZrO ball (Fig. 6A). Thereafter the MOF was washed with methanol, dried in a vacuum oven at 80°C for 36 h, and activated under ultrahigh vacuum at 120°C for 5 h. The material was characterized by PXRD (Fig. S14), thermogravimetric analysis (Figs. S15–16) and nitrogen sorption measurements, which revealed a Brunauer-Emmett-Teller (BET) surface area of 860 m^2^ g^−1^ (Fig. 6B, black), consistent with typical values for mechanochemically or solvothermally-prepared material (600-1100 m^2^ g^−1^) (*31,34*) (Fig. 6B, red).

**Fig. 6.**
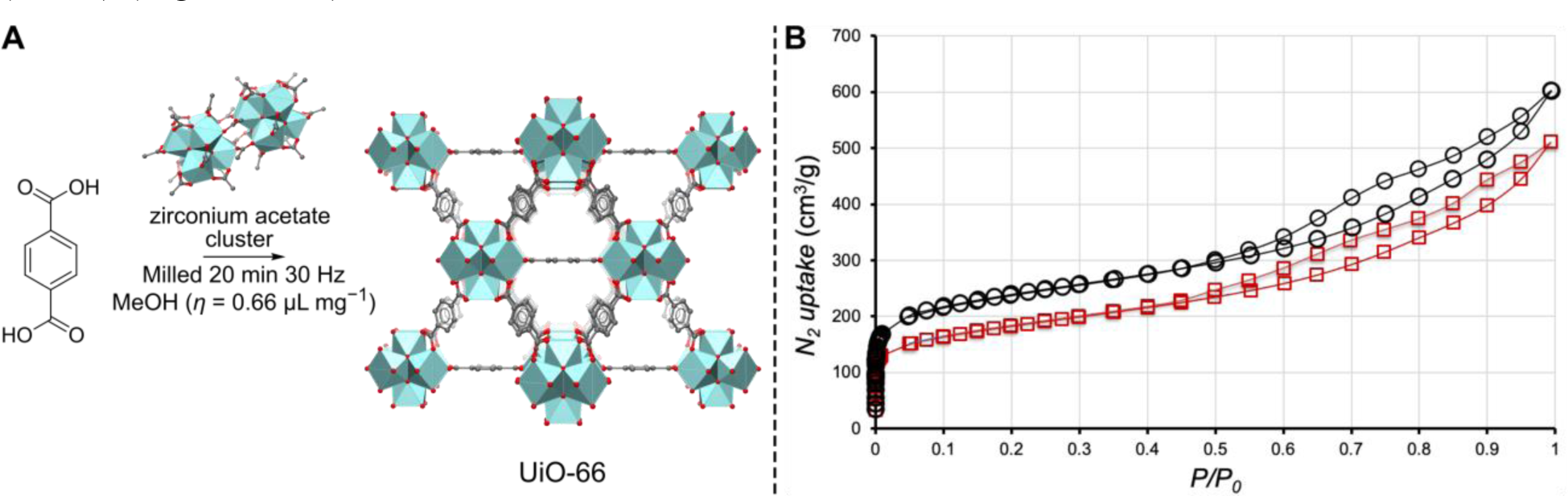
The UiO-66 MOF synthesized from TPA obtained by mechanoenzymatic PET hydrolysis. (A) The reaction scheme. (B) Overlay of nitrogen desorption and adsorption isotherms run at 77 K for UiO-66 synthesized either from herein recovered TPA (in black) or commercial TPA (in red) with surface area of 860 m^2^ g^−1^ and 640 m^2^ g^−1^, respectively.

## Conclusions

Using enzymes in moist-solid reaction mixtures rather than in traditional dilute solutions is an emergent strategy(*36*) that has proven beneficial for the breakdown of recalcitrant biopolymers such as cellulose,(*23, 24*) xylan,(*25*) and chitin(*26*). Such reaction conditions provide a milieu that is closer to the natural setting of enzymes like HiC, which are secreted in the environment by microorganisms. We demonstrate herein that, in contrast to enzymes used in dilute buffer, this approach enables the direct enzymatic depolymerization of high crystallinity PET commonly found in packaging, even when mixed with other plastics. Seven rounds of RAging produced >95% pure TPA in nearly 50% yield with only 3 wt% enzyme – by far the highest reported yield for highly crystalline PET – with a 13-fold higher STY and 15-fold higher yield per gram of enzyme compared to previous studies. This recycled TPA was found to be adequate for the manufacture of other value-added products, as illustrated herein with the production of the MOF UiO-66. Our results demonstrate that the high energy thermomechanical amorphization step, currently considered necessary for efficient enzymatic PET depolymerization(*12*), can be avoided simply by switching to moist-solid mechanoenzymatic reactions. Inherently minimalistic, our methodology proceeds with only the plastic substrate, the enzyme, a minor amount of buffer, and gentle, intermittent mechanical activation. Although validated here with the commercial wild type HiC enzyme, this method should be especially promising for engineered proteins. In addition to avoiding harsh chemicals, elevated temperatures, and high pressures, this strategy also minimizes the total reaction volume, thereby avoiding solubility issues, greatly facilitating handling and mixing, and curtailing waste. Overall, it enables the true recycling of PET plastics under milder, more sustainable conditions.

## Acknowledgments

We thank NSERC (RGPIN-2017-06467, SMFSU 507347-17), FRQNT (PR 254169), the McGill Sustainability Systems Initiative, and the Centre in Green Chemistry and Catalysis for funding (FRQNT-2020-RS4-265155-CCVC). We also thank Dr. Hatem Titi for his assistance with the nitrogen sorption measurements of the MOF materials, and Mr. Jean-Louis Do for providing the dodecanuclear zirconium acetate cluster.

## Author contributions

S.K. and J.P.D.T. conducted the research; D.D. contributed by the synthesis of the mono(2-hydroxyethyl) terephthalate standard; C.E. D. contributed with testing of the non-PET polymers; S.K. and J.P.D.T. wrote the original draft; K.A. and T.F. reviewed and edited the manuscript, in addition to supervising the research.

## Notes

### Competing Interest Statement

The authors have declared no competing interest.

