## Supplemental Methods, Figures and Tables for "Clean Enzymatic depolymerization of highly crystalline polyethylene terephthalate in moist-solid reaction mixtures"

**This PDF file includes:**

### Materials and Methods

Materials

Unless specified otherwise, water was from a MilliQ system with a specific resistance of 18.2 MΩ·cm at 25°C and all solvents used in product analysis (ACN, MeOH, DMSO, formic acid) were of HPLC grade, purchased from Millipore Sigma (Oakville, ON, Canada) and Thermo Fisher Scientific (Waltham, MA, US). Methanol (reagent grade, Thermo Fisher Scientific; Waltham, MA, US) and sodium carbonate (Thermo Fisher Scientific; Waltham, MA, US) aqueous solution were used for product extraction. NMR samples were prepared with $d_6$-DMSO and $d_4$-methanol, purchased from Cambridge Isotope Laboratories, Inc (Tewksbury, MA, US).

Sodium phosphate buffer was prepared from $NaH_2PO_4$ and $Na_2HPO_4$ (Chem-Impex; Wood Dale, IL, US), the Tris buffer was prepared from the Tris base (Chem-Impex; Wood Dale, IL, US) and HCl (Thermo Fisher Scientific; Waltham, MA, US). The bicine buffer was prepared from the bicine base (Millipore Sigma; Oakville, ON, Canada) and brought to pH 8 or 9 using HCl (Thermo Fisher Scientific; Waltham, MA, US).

The Novozym® 51032 enzyme used in this study was purchased from Strem Chemicals, Inc. (Newburyport, MA, US). Novozymes reports the source organism as *Humicola insolens*, with the enzyme expressed in an Aspergillus microorganism and is sold as a liquid solution. Unless noted otherwise, the commercial enzyme solution was used directly in reactions. To obtain lyophilized enzyme, the commercial HiC solution was dialyzed against 10 × 1 L of a 25 mM tris buffer (pH 7.3) containing 100 mM NaCl, and exchanged every hour for 10 hours. The HiC solution was first diluted to a concentration of 5 mg $mL^{-1}$ (from the initial 6.5 mg $mL^{-1}$) in the dialysis buffer before adding it (approximately 20 mL) to a Repligen Spectra/Por® 3 Standard Regenerated Cellulose (RC) Dialysis tube with a MWCO of 3.5 kDa (Spectrum Chemical Mfg. Corp.; New Brunswick, NJ, US). After dialysis, the content of the dialysis tube was placed into a 50 mL conical tube, frozen using liquid nitrogen, and lyophilized overnight using a Labconco (Kansas City, MO, US) FreeZone 1 L benchtop freeze-dry system. After lyophilization, the protein content of the solid powder and its activity towards BHET were measured.

The PET powder (particle size 300 μm) was purchased from Goodfellow (Coraopolis, PA, US). In addition, PET from a transparent post-consumer soft-drink bottle and an 80% recycled content black PET container were studied, both marked as polymer type 1. Post-consumer textile labelled as 100% polyester was kindly provided by Prof. Karine Auclair. All post-consumer materials were thoroughly washed with detergent, rinsed with MilliQ water, and dried in an oven at 55°C overnight. Polycarbonate (PC) granules, Bisphenol A, and polybutylene terephthalate (PBT) pellets were purchased from Millipore Sigma (Oakville, ON, Canada).

Standards for terephthalic acid (TPA), bis(2-hydroxyethyl) terephthalate (BHET) were purchased from Millipore Sigma (Oakville, ON, Canada). Mono-2-hydroxyethyl terephthalate (MHET) was synthesized from BHET according to the described method (See SI Methods: ***Preparation of Mono-2-Hydroxyethyl Terephthalate***). Benzoic acid, used as internal standard for NMR, was purchased from Sigma-Aldrich.

Methods

***Equipment***

Mettler Toledo AB135-S/FACT DualRange analytical balance (linearity 0.2 mg, readability 0.01 mg/0.1 mg), Mettler Toledo XP105 DeltaRange analytical balance (linearity 0.15 mg, readability 0.01 mg/0.1 mg), and Gilson and VWR pre-calibrated pipettors were used for measuring out the reaction components. Ball milling was carried out using a FormTech Scientific FTS1000 shaker mill, set at frequency 30 Hz, equipped with 15 or 30 mL sleeved PTFE or stainless-steel SmartSnap™ jars or unsleeved PTFE jars, all provided by FormTech Scientific, and charged either with $ZrO_2$ or stainless-steel balls. Aging steps of the experiments at 55°C were performed in a Fisher Scientific Oven, whereas aging at other temperatures and in-solution experiments were done using Shel Lab shaking incubators, VWR and Ohaus tabletop shaker incubators or Thermolyne Oven Series 9000. All reactions were performed in triplicate, unless noted otherwise. Error bars in graphs represent the standard deviations.

A Branson 2510 Sonicator was used to prepare samples for HPLC analysis, and Thermo Scientific Sorvall Legend Micro21 and Lynx 6000 centrifuges were used for centrifuging samples of up to 2 mL, and up to 25 mL, respectively. Lyophilization was carried out using a Labconco FreeZone 1 Liter Benchtop Freeze Dry System.

$^{1}H$ and $^{13}C$ solution-state NMR spectra were acquired on a Bruker AVIIIHD 500 MHz spectrometer. Chemical shifts are reported in ppm and referenced to the solvent residual signal (DMSO 2.50 ppm or MeOH 3.31 ppm). Spectra were analyzed and plotted using the TopSpin 4.0.6 software. Differential scanning calorimetry (DSC) measurements were conducted on a TA instruments Discovery 2500 Differential Scanning Calorimeter, on PET samples (2–8 mg in weight) placed into hermetically sealed aluminum pans. Powder X-ray diffraction analysis (PXRD) shown on **Fig. S4** was measured on a Bruker D8 ADVANCE X-ray diffractometer equipped with a Cu-Kα ($\lambda$ = 1.54 Å) source with a Ni filter and a LynxEye detector, in the $2\theta$ range of 10−40°, with step size of 0.02° and measuring time of 0.5 s per step. The PXRD patterns on **Fig. S14** were measured on a Bruker D2 Phaser diffractometer using nickel-filtered Cu-Kα radiation ($\lambda$ = 1.54184 Å), equipped with a LynxEye detector, in the $2\theta$ range of 4−40°, with step size of 0.025° and measuring time of 0.5 s per step. FTIR-ATR spectra were obtained in the 400 $cm^{-1}$ to 4000 $cm^{-1}$ range on a Bruker VERTEX 70 FTIR spectrometer with an integrated Platinum ATR unit. Scanning electron microscopy (SEM) images, with energy dispersive X-ray spectroscopy (EDX) analysis were collected on a FEI Quanta 450 Environmental Scanning Electron Microscope FE-ESEM.

Thermogravimetric analysis and differential scanning calorimetry data of the synthesized UiO-66 metal-organic framework were measured on a TGA/DSC 1 thermal analyzer (Mettler-Toledo, Ohio, USA) in open alumina crucibles, heated in the stream of air (25 mL $min^{-1}$) from 25 to 800°C at a heating rate of 10°C $min^{-1}$. Data collection and analysis were performed using the STARe Software 15.00 program package. The activation of the MOF materials was performed under ultrahigh vacuum at 120°C for 5 h on a Quantachrome instrument, before the surface area measurements. The porosity of the synthesized UiO-66 were determined by adsorption and desorption isotherm measurements performed on the Quantachrome instrument at a temperature of 77 K.

#### *Preparation of Mono-2-Hydroxyethyl Terephthalate (MHET)*

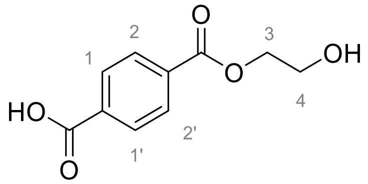

The reactant bis-2-hydroxy-ethyl terephthalate (BHET, 248 mg, 0.976 mmol) was added to 10 mL of 1:1 tetrahydrofuran:water. Lithium hydroxide monohydrate (46 mg, 1.10 mmol) was then added and the reaction mixture was stirred at room temperature for 16 hours. The reaction was then quenched with 2 M HCl to pH 2. The product was extracted into dichloromethane (1 × 10 mL). The organic layer was dried over sodium sulfate and the solvent was removed *in vacuo*. The solid was re-dissolved into a minimum of methanol for Prep-TLC (eluent: 1:9 methanol:dichloromethane). The streaked fraction with an $R_f$ between 0.2 and 0.6 was removed and the product was separated from the silica using a 1:9 methanol:dichloromethane mixture while filtering out the solid silica. The filtrate was dried to afford this know product as a white solid; 5 mg, 2%. $^1$H NMR (MeOD, 300 MHz) δ 8.17 – 8.09 (m, 4H, H-1, H-1', H-2, H-2'), 4.43 – 4.39 (m, 2H, H-3), 3.90 – 3.87 (m, 2H, H-4). These values correspond to the literature.(*1*)

#### *Protein characterization*

SDS-PAGE was performed with a 4–20% acrylamide gradient Mini-PROTEAN® TGX™ precast protein gel with 4 µL of the Amersham low-molecular weight ladder for SDS-PAGE or 10 µL of a 10-fold diluted commercial enzyme mix suspended in a 1× Laemmli buffer. The gel was stained with a Coomassie blue solution and imaged with a Bio-Rad ChemiDoc MP imaging system (Hercules, California, USA). The gels are shown on **Fig. S1**. The protein concentration was determined by using the commercial Thermo Scientific Pierce Coomassie Plus (Bradford) assay kit.

#### *Pre-milling conditions*

The post-consumer PET bottle, post-consumer black PET container (80% recycled content) and low crystallinity PET film were cut into roughly 0.7 × 0.7 cm squares and pre-milled at 30 Hz in a 30 mL stainless steel jar, using a 15 mm stainless steel ball (11.6 g) to obtain a powder. Milling 2 g of the bottle PET for 40 minutes yielded a powder with particle size distribution: 250 µm 10%, 250–500 µm 35%, 500–1000 µm 40% and >1000 µm 15%. The bottle PET powder with polypropylene impurity was obtained by milling the bottle squares with 10 wt% of cap pieces (red), resulting in a pink-colored powder. A powder containing 90% of particles below 1 mm was obtained from the post-consumer (80% recycled content) black PET container by milling for 30 minutes. The low crystallinity PET film was milled in installments of 10 min + 5 min + 5 min + 10 min, to allow the jar to return to room temperature. The obtained powder contained more flakes of >1 mm size than the powdered post-consumer PET materials and had **not** retained its low crystallinity. The crystallinity of all PET materials used was determined by differential scanning calorimetry (**Figs. S2**, **S3**).

The PC flakes were pre-milled by adding 2 g of the plastic to a 30 mL stainless steel jars containing a 15 mm stainless steel ball (11.6 g). The plastic was milled at 30 Hz for 30 minutes to obtain a powder.

The PBT pellets were pre-milled by adding 1 g of the plastic to a 30 mL stainless steel jars containing a 15 mm stainless steel ball (11.6 g). The plastic was milled at 30 Hz for 2 h to obtain a powder. After pre-milling, the powder was sieved to remove any particles larger than 1 mm.

***Crystallinity determination of PET materials according to DSC***

The degree of crystallinity ($X_c$) of the semi-crystalline PET polymers were estimated using the TRIOS software version 4.5.1.42498, based on heating scans from 0°C to 300°C at a rate of 10°C min$^{-1}$, according to the equation $X_c = \frac{\Delta H_f - \Delta H_c}{\Delta H_f^o}$, where $\Delta H_f$ is the enthalpy of fusion, $\Delta H_c$ is the enthalpy of cold crystallization (only exhibited by low crystallinity Goodfellow PET film) and $\Delta H_f^o$ is the enthalpy of fusion of a 100% crystalline PET polymer (140 J g$^{-1}$) at the melting temperature. Both the endothermic melting and the exothermic cold crystallization peaks were integrated from visually determined respective starting points to end points using a straight virtual baseline between them to get $\Delta H_f$ and $\Delta H_c$. The degree of crystallinity of the PET samples may be slightly overestimated, as they are not corrected for the temperature dependence of the melting enthalpy $\Delta H_f^o$.(*2*)

***Method for the mechanochemical + aging reactions***

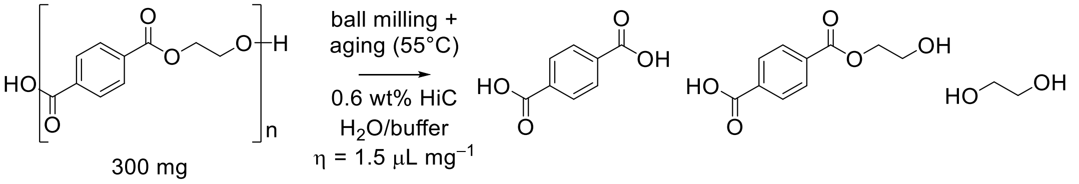

(Scheme S1)

In a typical reaction, PET (300 mg, 1.58 mmol) was weighed into a 15 mL PTFE or stainless-steel jar, charged either with $ZrO_2$ or stainless-steel ball(s), to which the commercial enzyme preparation (300 μL, 1.95 mg protein) and buffer (150 μL) were added. The liquid-to-solid ratio is defined as $\eta$ and expressed in μL mg$^{-1}$. The jar was then set to mill at a set frequency (generally 30 Hz) for 5 minutes. The resulting solids were aliquoted into three parts, 10–50 mg each, for analysis of the reaction products by HPLC at three time points: after milling, after 3 days of aging at 55°C, and after 7 days of aging at 55°C. Separate aliquots were prepared for PXRD and NMR analysis of the reaction products, and analyzed after 3 or 7 days of aging at 55°C. Specific reaction conditions and variations tested, together with corresponding hydrolysis yields are compiled into **Table S1 and Table S3**.

***Method for the mechanochemical + aging reactions using the lyophilized enzyme***

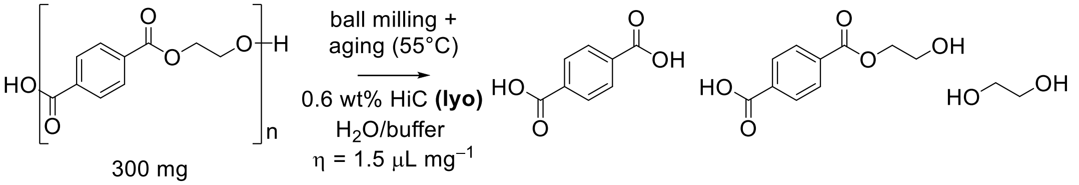

(Scheme S2)

PET (300 mg, 1.58 mmol) was weighed into a 15 mL stainless-steel jar, charged with a single 9 mm stainless-steel ball (3.5 g), to which the lyophilized enzyme powder (16.3 mg, 1.95 mg protein) and buffer/water (150–450 μL) were added. The liquid-to-solid ratio is defined as $\eta$ and expressed in μL mg$^{-1}$. The jar was then set to mill at a set frequency (30 Hz) for 5 minutes. The resulting solids were aliquoted into three parts, 10–50 mg each, for analysis of the reaction products by

HPLC at three time points: after milling, after 3 days of aging at 55°C, and after 7 days of aging at 55°C. Separate aliquots were prepared for PXRD and NMR analysis of the reaction products, and analyzed after 3 or 7 days of aging at 55°C. Specific reaction conditions and variations tested, together with corresponding hydrolysis yields are compiled into **Table S2**.

#### *Studying the influence of non-participating polymers on the mechanoenzymatic reactions*

PET (100 mg) and the non-participating polymer (200 mg) were weighed into a 15 mL stainless steel jar, charged with a single 9 mm stainless-steel ball (3.5 g), to which the commercial enzyme preparation (300 μL, 1.95 mg protein) and 0.1 M sodium phosphate buffer at pH 7.3 (150 μL) were added. The liquid-to-solid ratio $\eta$ was 1.5 μL mg$^{-1}$. The jar was then set to mill at 30 Hz for 5 minutes.

The resulting solids after milling were split into 10–50 mg aliquots for the analysis of reaction products by HPLC at three time points: after milling, after 3 days of aging at 55°C, and after 7 days of aging at 55°C. Specific reaction conditions and variations tested, together with corresponding hydrolysis yields are compiled into **Table S4**.

#### *Mechanoenzymatic depolymerization of other plastic polymers*

In a typical reaction, PBT, PC, or PCL (300 mg) were weighed into a 15 mL stainless-steel jar, charged with a 9 mm stainless-steel ball (3.5 g), to which the commercial enzyme preparation (300 μL, 1.95 mg protein) and buffer (150 μL) were added. The jar was then set to mill at a set frequency (30 Hz) for 5 minutes. The resulting solids were aliquoted into three parts, 10–50 mg each, for analysis of the reaction products by HPLC (PBT and PC) or NMR (PCL) at three time points: after milling, after 3 days of aging at 55°C, and after 7 days of aging at 55°C. Specific reaction conditions and variations tested, together with corresponding hydrolysis yields are compiled into **Table S5**.

#### *Method for the Reactive Aging reactions (RAging)*

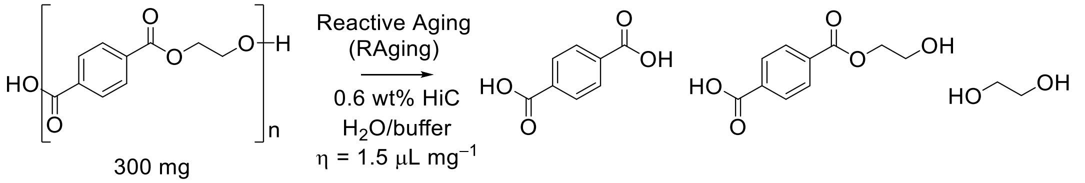

(Scheme S3)

In a typical RAging reaction, PET (300 mg, 1.58 mmol) was weighed into a 15 mL unsleeved PTFE jar, charged with a single 10 mm $ZrO_2$ ball (3.5 g), to which the commercial enzyme preparation (300 μL, 1.95 mg protein) and buffer (150 μL) were added. A cycle of RAging typically consisted of: ball milling at 30 Hz for 5 minutes followed by aging at 55°C for 24 hours. These cycles were repeated up to 7 times, with an aliquot (10–20 mg) collected at each cycle for analysis of the reaction products by HPLC. Sealing tape proved to be necessary to ensure that the jars remain closed during the aging step. Specific reaction conditions and variations tested, together with corresponding hydrolysis yields are compiled into **Table S6**.

#### *Method for the product inhibition studies*

**Terephthalic acid (TPA) inhibition**: PET (250 mg, 1.30 mmol) and TPA (50 mg, 0.3 mmol) were weighed into a 15 mL stainless steel jar, charged with a single 9 mm stainless-steel ball (3.5 g), to which the commercial enzyme preparation (300 μL, 1.95 mg protein) and 0.1 M sodium phosphate buffer pH 7.3 (150 μL) were added. The liquid-to-solid ratio $\eta$ was 1.5 μL $mg^{-1}$. The jar was then set to mill at 30 Hz for 5 minutes.

**Terephthalic acid (TPA) and ethylene glycol (EG) inhibition**: PET (250 mg, 1.30 mmol) and TPA (50 mg, 0.3 mmol) were weighed into a 15 mL stainless steel jar, charged with a single 9 mm stainless-steel ball (3.5 g), to which EG (20 μL) was added. The jar was closed and milled for 1 minute, to mix the liquid EG well with PET and TPA. After that, the commercial enzyme preparation (300 μL, 1.95 mg protein) and 0.1 M sodium phosphate buffer pH 7.3 (150 μL) were added. The liquid-to-solid ratio $\eta$ was 1.57 μL $mg^{-1}$ when counting in EG. The jar was then set to mill at 30 Hz for 5 minutes.

**Ethylene glycol (EG) inhibition**: PET (300 mg, 1.58 mmol) was weighed into a 15 mL stainless steel jar, charged with a single 9 mm stainless-steel ball (3.5 g), to which EG (150 μL) was added. The jar was closed and milled for 10 minutes, to mix the liquid EG well with PET. After that, the commercial enzyme preparation (300 μL, 1.95 mg protein) and water (150 μL) were added. A control reaction was carried out by adding only water (450 μL) and no enzyme. The liquid-to-solid ratio $\eta$ was 2 μL $mg^{-1}$ when counting in EG. The jar was then set to mill at 30 Hz for 5 minutes.

For all three conditions, the resulting solid after milling was split into 10–50 mg aliquots for the analysis of reaction products by HPLC at three time points: after milling, after 3 days of aging at 55°C, and after 7 days of aging at 55°C. Specific reaction conditions and variations tested, together with corresponding hydrolysis yields are compiled into **Table S7**.

#### *HPLC separation and characterization of PET and PBT hydrolysis products*

The hydrolysis products from a typical PET hydrolysis reaction after aging were extracted by adding a corresponding amount of DMSO to the reaction aliquot (to a 20 or 25 mg $mL^{-1}$ solution). The dissolution of all hydrolysis products from the reaction mixture was ensured by sonicating the mixture for 1–5 minutes at room temperature. The suspension was thereafter centrifuged at 21 000 × *g* for 5 minutes to separate the remaining PET from the solution. An appropriate volume of the supernatant was diluted 10- to 40-fold in MeOH to prepare for HPLC analysis. The hydrolysis products from the PBT reactions were prepared at 20 mg $mL^{-1}$ concentration in MeOH without any further dilution. All samples were syringe filtered using a 0.22 µm nylon filter (Chromspec, Brockville, ON, Canada) to remove any fine particles prior to HPLC analysis. Samples from reaction mixtures straight after milling (at low TPA conversion) were suspended in MeOH to a concentration of 20 mg $mL^{-1}$ solids, with the supernatant directly analyzed by HPLC after sonication, centrifugation and filtration.

Hydrolysis products of PET and PBT were quantified using HPLC equipped with a quaternary pump, autosampler, and multiwavelength UV–vis detector (1260 Infinity II, Agilent Technologies, Santa Clara, CA, USA). Separation was achieved on a Phenomenex Luna® 5 µm C18(2) 100 Å, 4.6 × 250 mm column using a 40-minute gradient elution from 99(A):1(B) to 60(A):40(B), where A is 0.1% formic acid in water and B is acetonitrile. The injection volume was set to 10 μL, the flow rate used was 0.6 mL $min^{-1}$ and UV detection was performed at 240 nm.

TPA, MHET, and BHET standards were prepared at a 10 mg $mL^{-1}$ stock in DMSO and were diluted to 200, 100, 50, and 25 µg $mL^{-1}$ in MeOH. Samples were syringe filtered using a 0.22 µm nylon filter (Chromspec, Brockville, ON, Canada) to remove any fine particles. The areas under the peaks were used to construct the calibration curves, which were then used to calculate the concentration of TPA, MHET and BHET in the samples.

***Method of calculating the yield of TPA (% of theoretical)***

The percent TPA yields reported in this paper are calculated by dividing the experimentally determined yield of TPA (mg), extrapolated from the content of TPA in solids as determined by the HPLC analysis, with the theoretical yield at 100% conversion. Assuming a 100% pure polyethylene terephthalate material, complete hydrolysis of for example 300 mg (1.58 mmol) of PET would yield 259.3 mg (1.58 mmol) of TPA and 96.9 mg (1.58 mmol) of ethylene glycol.

***Optimization of TPA isolation***

Three isolation methods for the hydrolysis products were tested on bulk reaction mixtures at 18–20% conversion to TPA.

*Method A.* Hydrolysis products were extracted from the solid reaction mixture (from 300 mg PET) with 11 × 10 mL methanol, which were combined and dried *in vacuo*. $^{1}$H NMR analysis (**Fig. S11**) showed that the product thus extracted (120 mg) contains a significant amount of 1,2-propylene glycol (5 molar equivalents to TPA), the cryoprotectant present in the commercial enzyme preparation. The amount of TPA obtained was 33 mg, yield 13%.

*Method B.* The solid reaction mixture (from 300 mg PET) was first washed with 3 × 1 mL of MilliQ water to remove the cryoprotectant and other water-soluble reaction components, after which TPA was extracted in 11 × 10 mL methanol which were combined and dried *in vacuo*. The product thus isolated (42.5 mg) contains mainly TPA (91.5 wt%) according to $^{1}$H NMR analysis (**Fig. S12**), with the minor impurity being MHET (8.5 wt%). Therefore, the amount of TPA obtained was 39 mg, yield 15%.

*Method C.* The TPA in the solid reaction mixture (from 900 mg PET) was reacted with 1.2 eq of sodium carbonate (added as 0.15 M aqueous solution), to yield a water-soluble sodium terephthalate salt that was easily extracted from the remaining PET with 3 × 3 mL MilliQ water. The combined aqueous solutions of sodium terephthalate were filtered through cotton and acidified with 2 molar equivalents of HCl to pH 1–2, resulting in the precipitation of TPA. The TPA was isolated by centrifugation (5000 × *g*) and removal of the acidic supernatant. Ideally, higher centrifugal forces should be used, as the supernatant removed was not fully clear and some product was thus lost. The product was washed with water until washings were neutral, and lyophilized to yield a white powder (98 mg). According to $^{1}$H NMR analysis (**Fig. S13**) the isolated product contains mainly TPA (96 wt%), with the minor impurity from MHET (4 wt%). Therefore, 94 mg of TPA was isolated, yield 12%.

The isolated reaction products were also analyzed by PXRD (**Fig. S4**) and FTIR (**Fig. S6**).

***Determining the kinetics of aging reactions***

To determine the reaction kinetics of aging reactions, PET (300 mg, 1.58 mmol) was weighed into a 15 mL stainless-steel jar, charged with a 9 mm (3.5 g) stainless-steel ball, to which the commercial enzyme preparation (300 µL, 1.95 mg protein) and 0.1 M sodium phosphate buffer at pH 7.3 (150 µL) were added. The jar was then set to mill at 30 Hz for 5 minutes. The resulting solids were aliquoted into 16 parts, 10–50 mg each, for the analysis of reaction products by HPLC at 16 time points: after milling, then after 30 min, 1 h, 2 h, 3 h, 4 h, 5 h, 6 h, 8 h, 10 h, 12 h, 16 h, 24 h, 48 h, 72 h, and 96 h of aging at 55°C.

#### *In-solution BHET enzyme activity assay*

The activity of HiC in the commercial Novozym® 51032 preparation and in the washings of the solid reaction mixtures was determined by an enzyme assay following the hydrolysis of bis(2-hydroxyethyl) terephthalate (BHET). This assay was carried out using solutions containing 0.65 mg $mL^{-1}$ enzyme, diluted in buffer (0.1 M sodium phosphate buffer at pH 7.3) as necessary. Namely, the enzyme solution (99 μL) was incubated at 55°C for 10 minutes, after which the reaction was initiated with the addition of 1 μL of a 100 mM BHET stock in DMSO, to achieve a final concentration of 1 mM BHET, and allowed to shake (250 rpm) at 55°C. The reaction was quenched by adding MeOH (400 μL), followed by sonication and centrifugation, after which the clear supernatant was analyzed by HPLC. HiC in the commercial preparation converted ~99% of the BHET into MHET after only 10 min, with no loss in enzymatic activity over several months. The results of the enzyme activity assays are compiled into **Table S8**.

#### *In-solution BHET enzyme activity assay on the HiC adsorbed on the remaining hydrolyzed PET solids*

Dried remaining PET solid (10 mg), washed with either methanol (see ***TPA isolation*** *Method A*) or 1 M Tris-HCl pH 8.2 buffer (see ***TPA isolation*** *Method* B) and dried by lyophilization, was added to 100 μL of 1 mM solution of BHET in 0.1 M sodium phosphate buffer at pH 7.3, and allowed to shake at 250 rpm for 2 hours at 55°C. The reaction was quenched by addition of MeOH (400 μL), followed by sonication and centrifugation, after which the clear supernatant was analyzed by HPLC. Each solid was tested in triplicate, and the consumption of BHET was monitored. The results of the enzyme activity assays are compiled into **Table S8**.

#### *Multi-round RAging experiments*

PET (900 mg, 4.74 mmol) was weighed into a 15 mL unsleeved PTFE jar, charged with a single 10 mm $ZrO_2$ ball (3.5 g), to which the buffer (450 μL) and the commercial enzyme preparation (900 μL, 5.85 mg protein) were added. Control reactions were carried out with the flow-through (900 μL) from the commercial enzyme solution, obtained by using a 10 kDa MWCO centrifugal concentrator. Sealing tape proved to be necessary to ensure that the jars remain closed during the aging steps. The multi-round experiment consisted of seven rounds, three RAging cycles each (5 minutes of milling at 30 Hz, followed by aging at 55°C for 24 hours) – totaling 21 RAging cycles. After every round, hydrolysis products were removed by transferring the solid paste to a 15 mL conical tube, extracting TPA in 0.3 M $Na_2CO_3$, vortexing and centrifuging at 10,000 × *g* for 5 min. The supernatant containing the soluble sodium terephthalate was removed and TPA was recovered as described in *Method C* under ***TPA isolation***. The remaining PET solid was washed with MilliQ water (3 × 3 mL), until neutral pH was reached, frozen with liquid nitrogen and lyophilized. The remaining PET was then used for the next round: weighed into a 15 mL unsleeved PTFE jar, charged with a single 10 mm $ZrO_2$ ball (3.5 g), and the corresponding amount of enzyme preparation (0.65% $w_{protein}/w_{solid}$) and buffer to maintain an $\eta$ of 1.5 μL $mg^{-1}$. An aliquot (10–20 mg) was taken for HPLC analysis at the start and end of each round. The results are displayed on **Fig. 5** (main text). The remaining PET was characterized by SEM (with EDX) after $1^{st}$ and after $7^{th}$ round, as seen in **Figs. S9** and **S10**, and compared with non-hydrolyzed PET (**Fig. S8**).

#### *In-solution control of enzymatic depolymerizations*

PET, PBT, or PC (20 mg $mL^{-1}$ ) was suspended in 0.1 M sodium phosphate buffer at pH 7.3 in the presence of 0.65% protein ($w_{protein}/w_{solid}$). Samples were placed in a shaking incubator set at 250 rpm for up to 7 days at 55°C. After 3 or 7 days, samples were frozen with liquid nitrogen and lyophilized. The solid was then analyzed by HPLC as described previously. The hydrolysis yields are shown in **Table S1**, **line 24** (PET) and **Table S5 lines 4–6** (PBT, PC, PCL).

#### *Synthesis of the UiO-66 metal-organic framework (MOF)*

The MOF materials were synthesized analogously to previously reported mechanochemical methods (refs. 31–33 in the main text), from zirconium acetate cluster $Zr_{12}O_8(OH)_8(CH_3COO)_{24}$ and either from the isolated TPA (isolated according to *method C*, see SI Methods: ***TPA isolation***) or from the purchased reagent grade TPA. The zirconium acetate cluster was synthesized according to literature procedure.(*3*)

TPA (52.5 mg, 0.32 mmol) and the zirconium acetate cluster (90 mg, 0.032) were weighed into a 15 mL $ZrO_2$ jar, charged with a 3.2 g $ZrO_2$ ball, to which 94 µL of methanol ($\eta$ = 0.66 µL $mg^{-1}$) was added. The jar was closed and set to mill for 20 minutes at 30 Hz. The resulting caking powder was transferred to an Eppendorf and washed with methanol. First the solids were washed with 3 × 1 mL, centrifuging the solids between washes at 21 000 × *g* for 3 minutes. Then the solids were resuspended in 1 mL of methanol and allowed to soak overnight (16 h). After removing the supernatant, the solids were again washed with 3 × 1 mL of methanol and then set to dry in glass vials in a vacuum oven at 80°C for 36 h. The obtained powders were characterized by PXRD (**Fig. S14**) and TG/DSC (**Figs. 15** and **16**), then activated under ultrahigh vacuum at 120°C for 5 h, prior to measuring the nitrogen adsorption-desorption isotherms (shown on **Fig. 6B** in the main text).

#### *HPLC separation and characterization of HiC-catalyzed PC depolymerization products*

Hydrolysis products of PC were quantified using HPLC with quaternary pump, autosampler, and multiwavelength UV–vis detector (1260 Infinity II, Agilent Technologies, Santa Clara, CA, USA). Separation was achieved with a ZORBAX Extend-C18, 5 µm, 80 Å, 4.6 × 250 mm column using a water:acetonitrile isocratic elution of 60:40 for 7 min at a flow rate of 0.6 mL/min. UV detection was performed at 260 nm and 280 nm.

BPA standards were prepared as 10 mg $mL^{-1}$ stock solutions in MeOH, diluted to 5.0, 1.0, 0.5, and 0.1 mg $mL^{-1}$ in MeOH. Samples were syringe filtered using a 0.22 µm nylon filter (Chromspec, Brockville, ON, Canada) to remove any fine particles. The area under the peak was used to construct the calibration curve, which was then used to calculate the concentration of the samples.

### Supplementary Figures

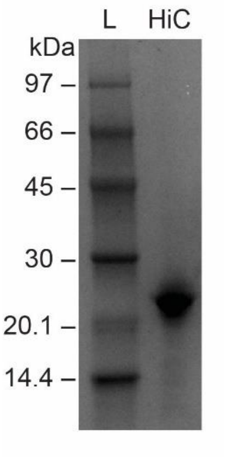

**Fig. S1.**

Characterization of the commercial *Humicola insolens* cutinase (HiC) solution marketed as Novozym® 51032. The commercial solution was analyzed using SDS-PAGE with a 4–20% acrylamide gradient gel stained with Coomassie blue.

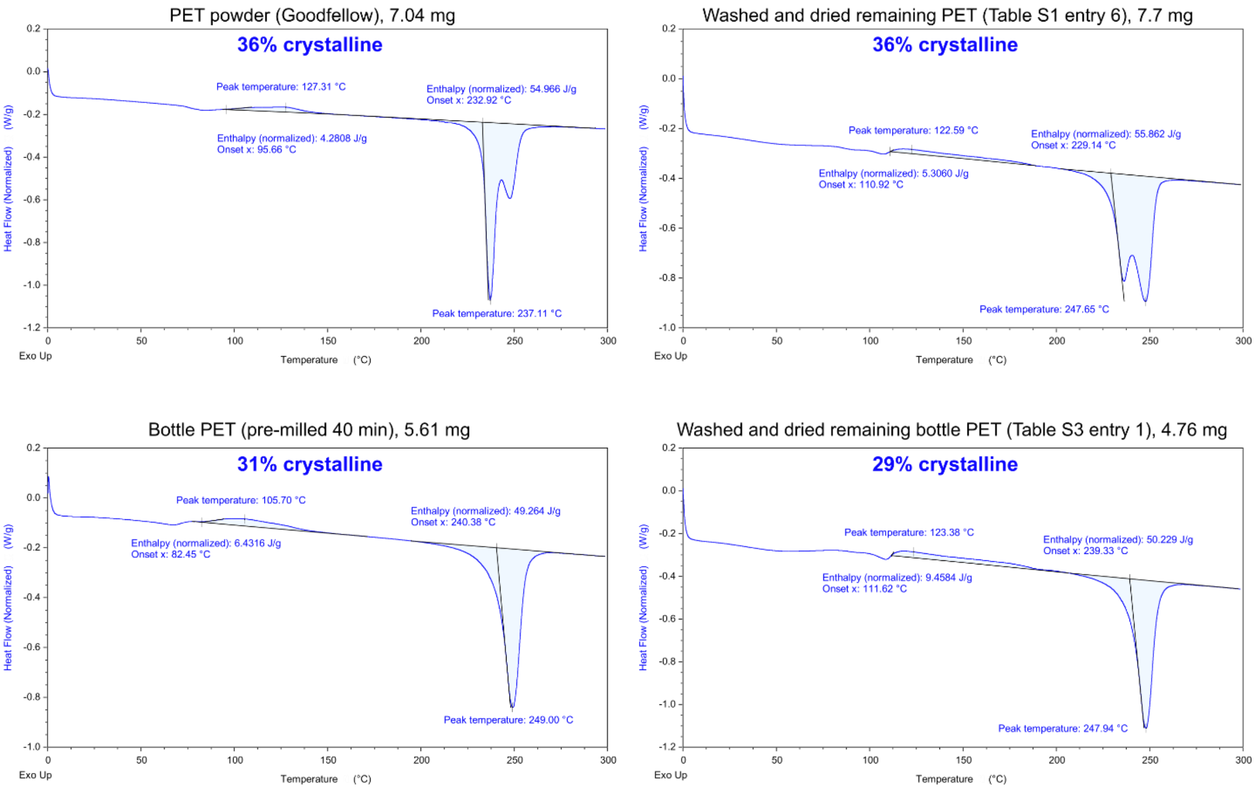

**Fig. S2.**

Heating scans from 0°C to 300°C at a rate of 10°C min$^{-1}$ on PET powder (Goodfellow) and bottle PET (pre-milled 40 minutes), showing the crystallinity the PET materials before (on the left) and after (on the right) the reaction according to the DSC analysis.

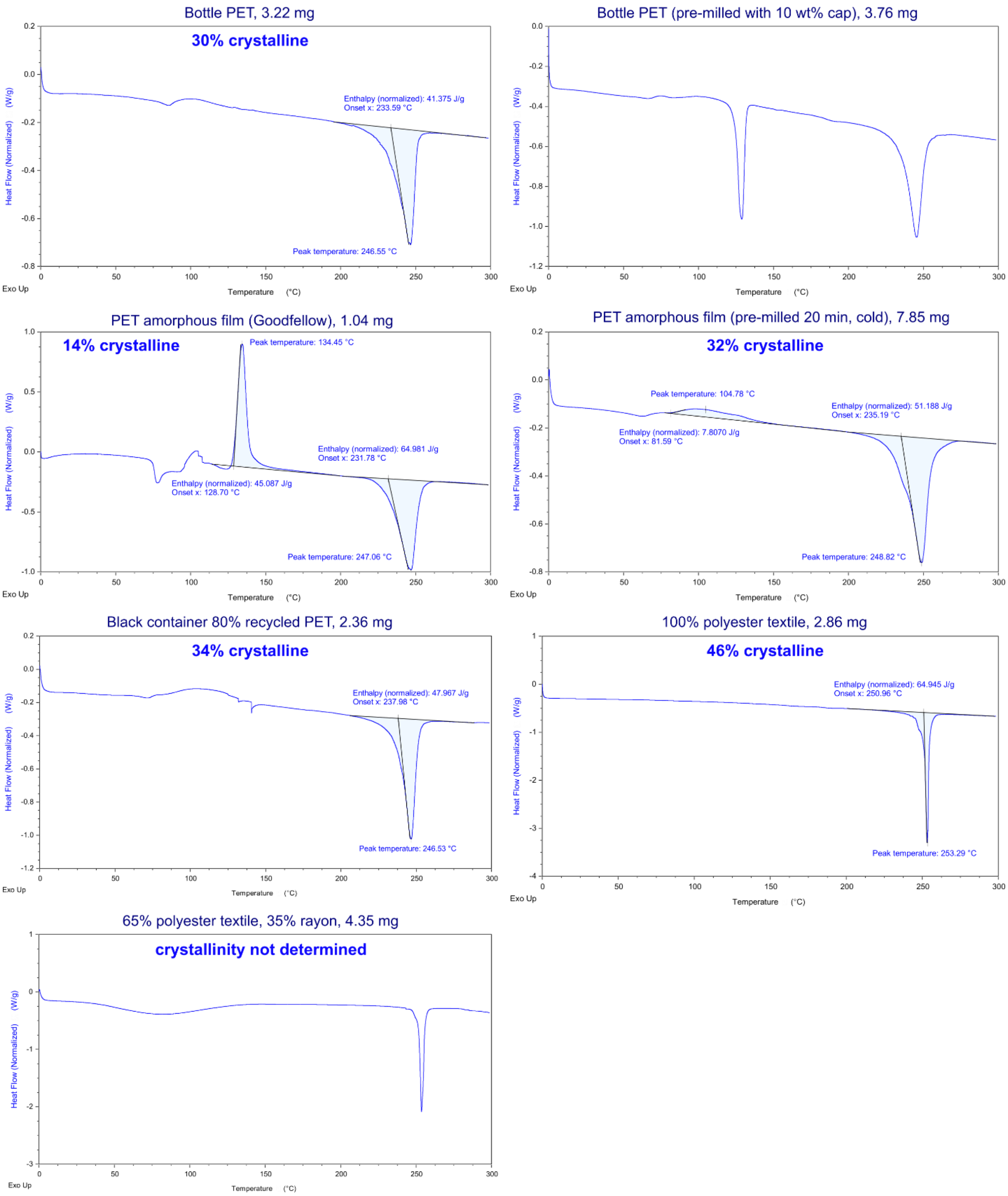

**Fig. S3.**

Heating scans from 0°C to 300°C at a rate of 10°C min$^{-1}$ on bottle PET powder, bottle PET powder with 10 wt% of the cap material (polypropylene, mp 130°C), low-crystallinity thin film before and after pre-milling, black PET container (80% recycled content), and textiles labelled as 100% polyester, and 65% polyester / 35% rayon. The crystallinity the PET materials according to the DSC analysis are shown on the scans. PET crystallinity in mixed polymers was not determined.

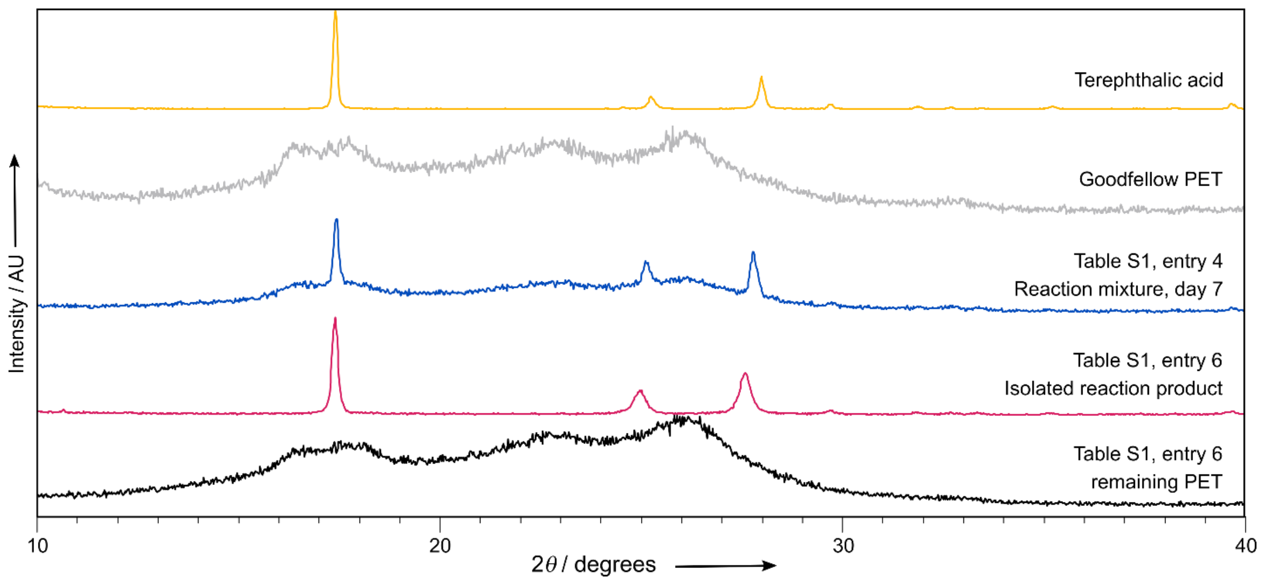

**Fig. S4.**

The Powder X-ray diffraction (PXRD) patterns of terephthalic acid (in yellow) corresponding to the crystal structure CCDC 1269122 from ref. (*4*); the Goodfellow PET powder (in gray) as purchased; a reaction mixture at day 7 (in blue) showing that the crude reaction product contains crystalline TPA; the powder patterns of the isolated reaction product (in magenta, isolated according to method A, see SI Methods: ***TPA isolation***), which corresponds in full to the powder pattern of TPA; and the remaining PET (in black), after the products were isolated, showing that the PET has retained the same PXRD features as PET before hydrolysis.

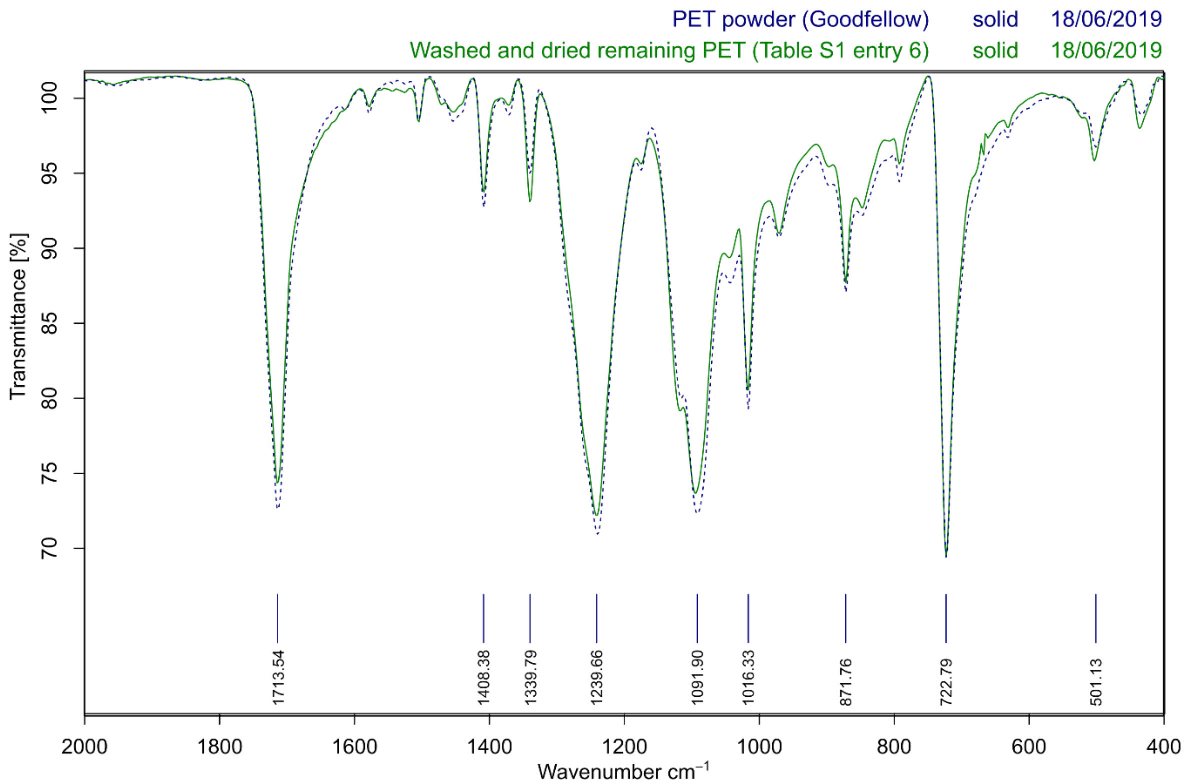

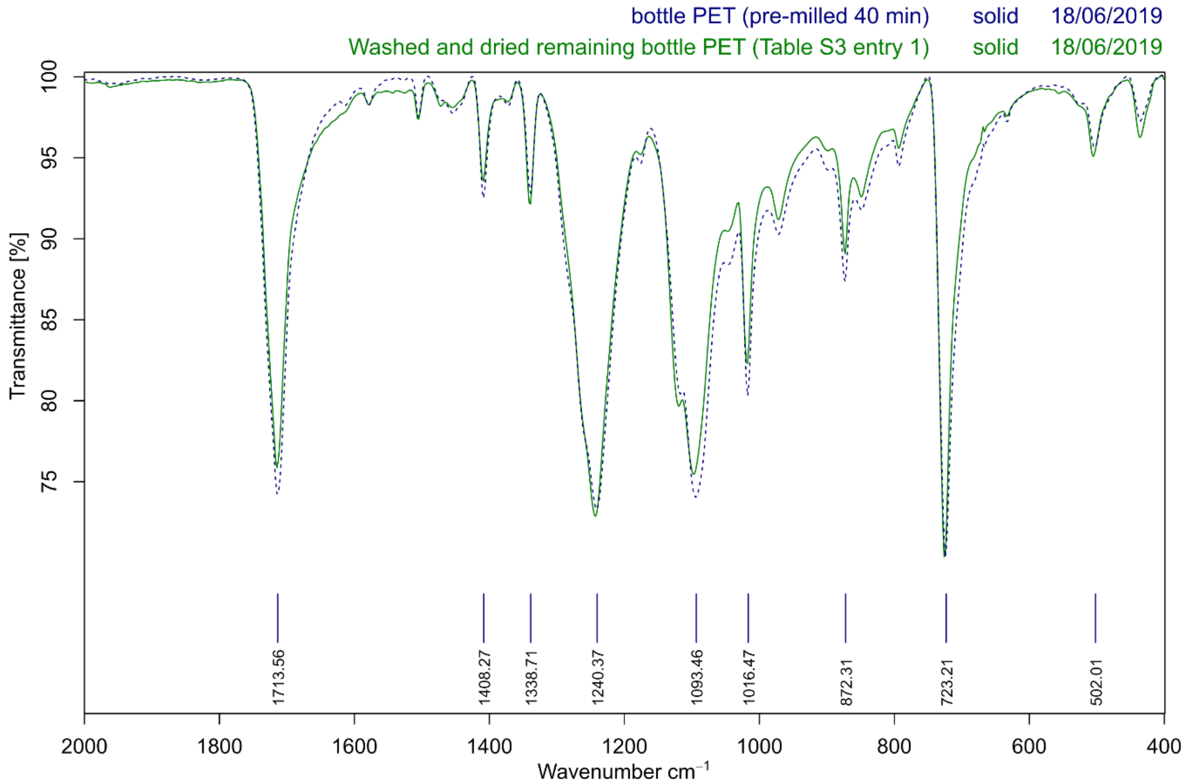

**Fig. S5.**

FTIR spectra of PET (powder and bottle) before and after the mechanoenzymatic reaction. Only the range 400–2000 cm$^{-1}$ is shown for clarity.

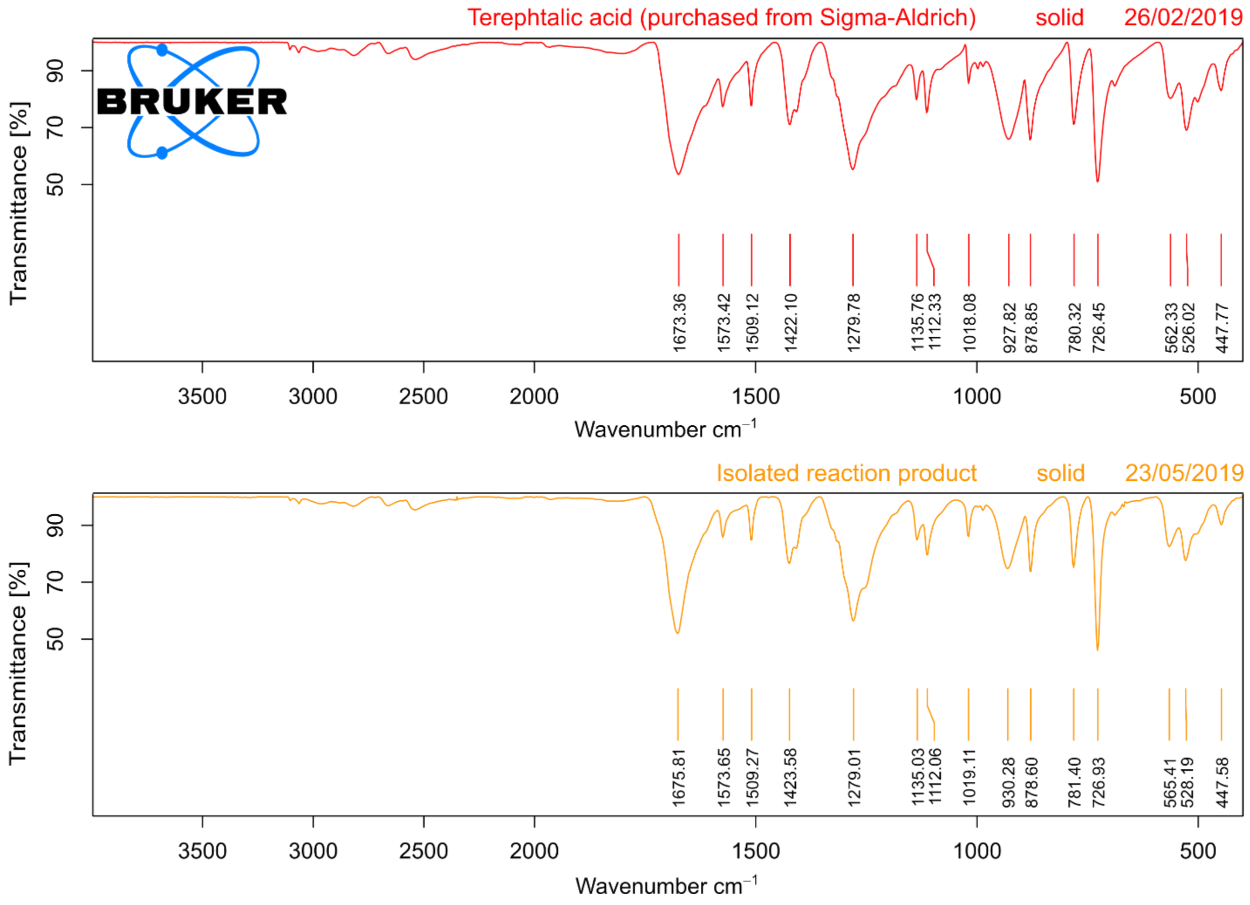

**Fig. S6.**

FTIR spectra of the isolated reaction product (in yellow, isolated according to method A, see SI Methods: ***TPA isolation***) in comparison with a commercially sourced TPA (red).

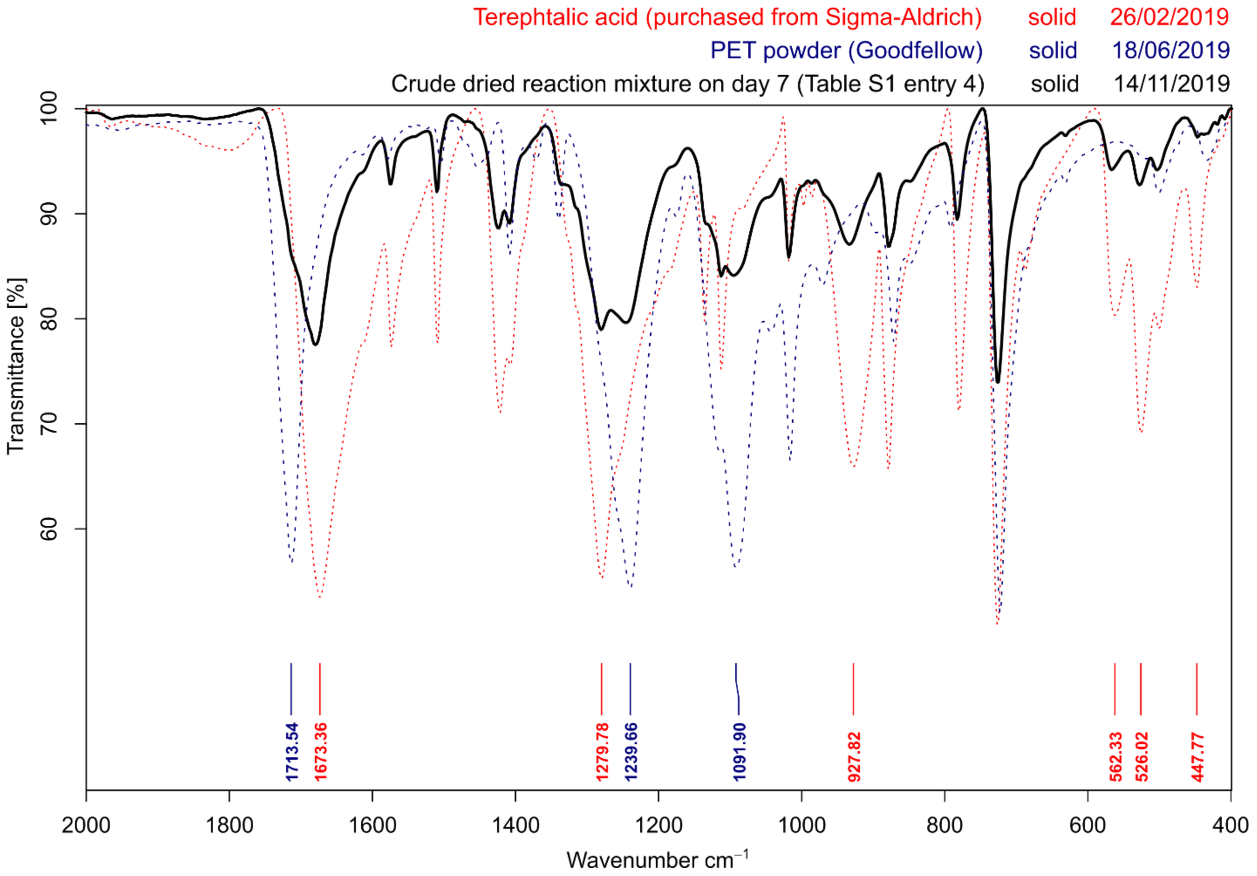

**Fig. S7.**

FTIR spectra of the starting material (in blue dotted line, PET powder from Goodfellow), a crude dried reaction mixture at day 7 (in black solid line) and the expected reaction product TPA (in red dotted line). Wavenumbers of peaks characteristic for TPA (red) and PET (blue), used to gauge the composition of the crude reaction mixture after aging. Only the range 400–2000 cm$^{-1}$ is shown for clarity.

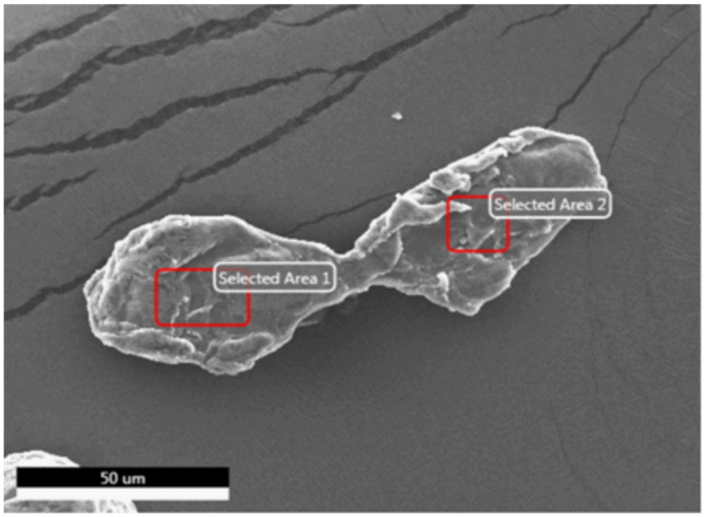

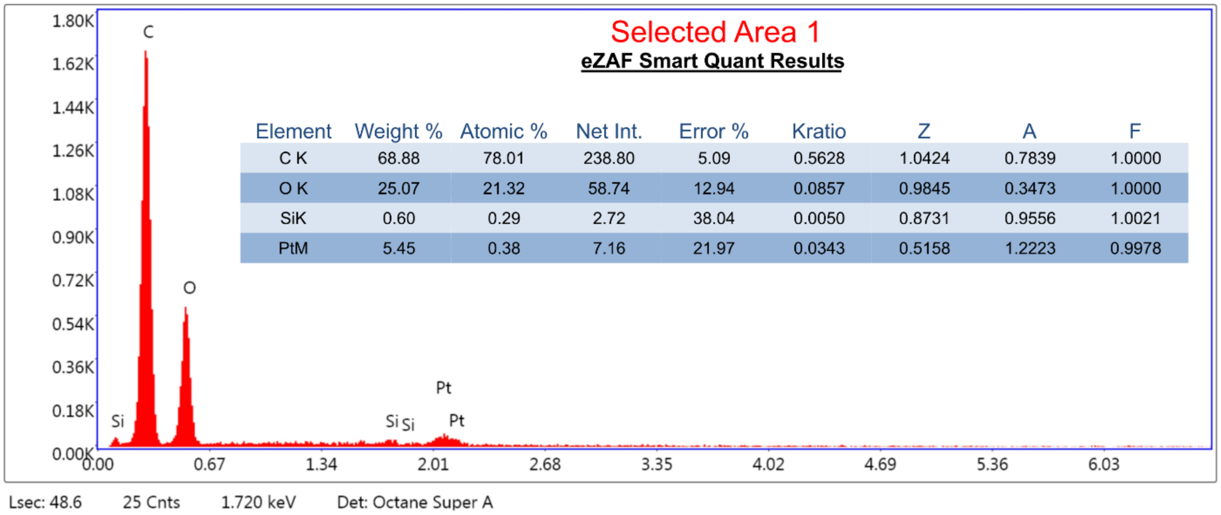

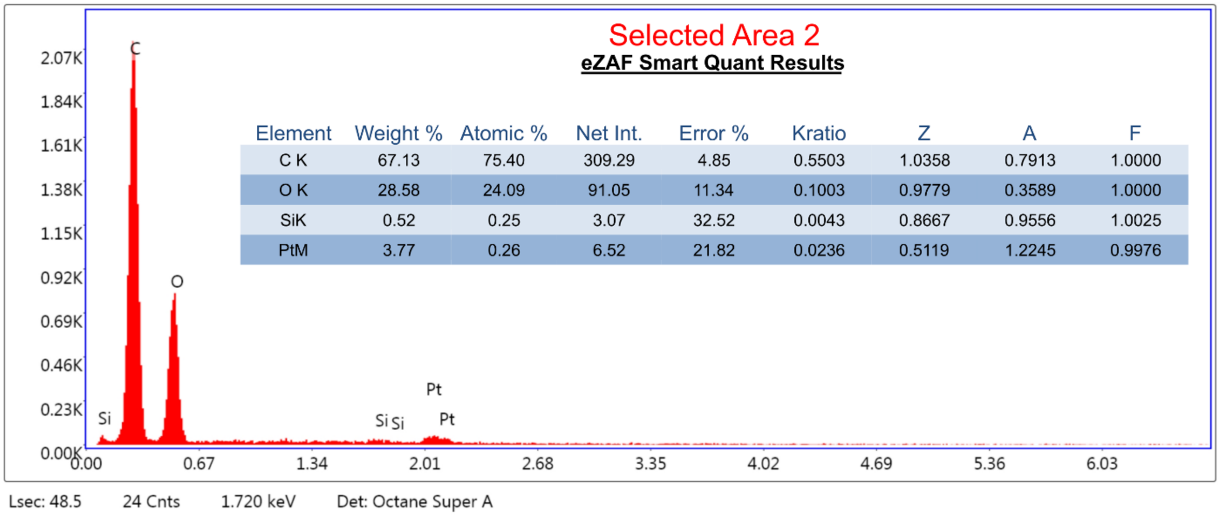

**Fig. S8.**

SEM image of the PET material before hydrolysis, and the corresponding EDX spectra indicating the absence of nitrogen.

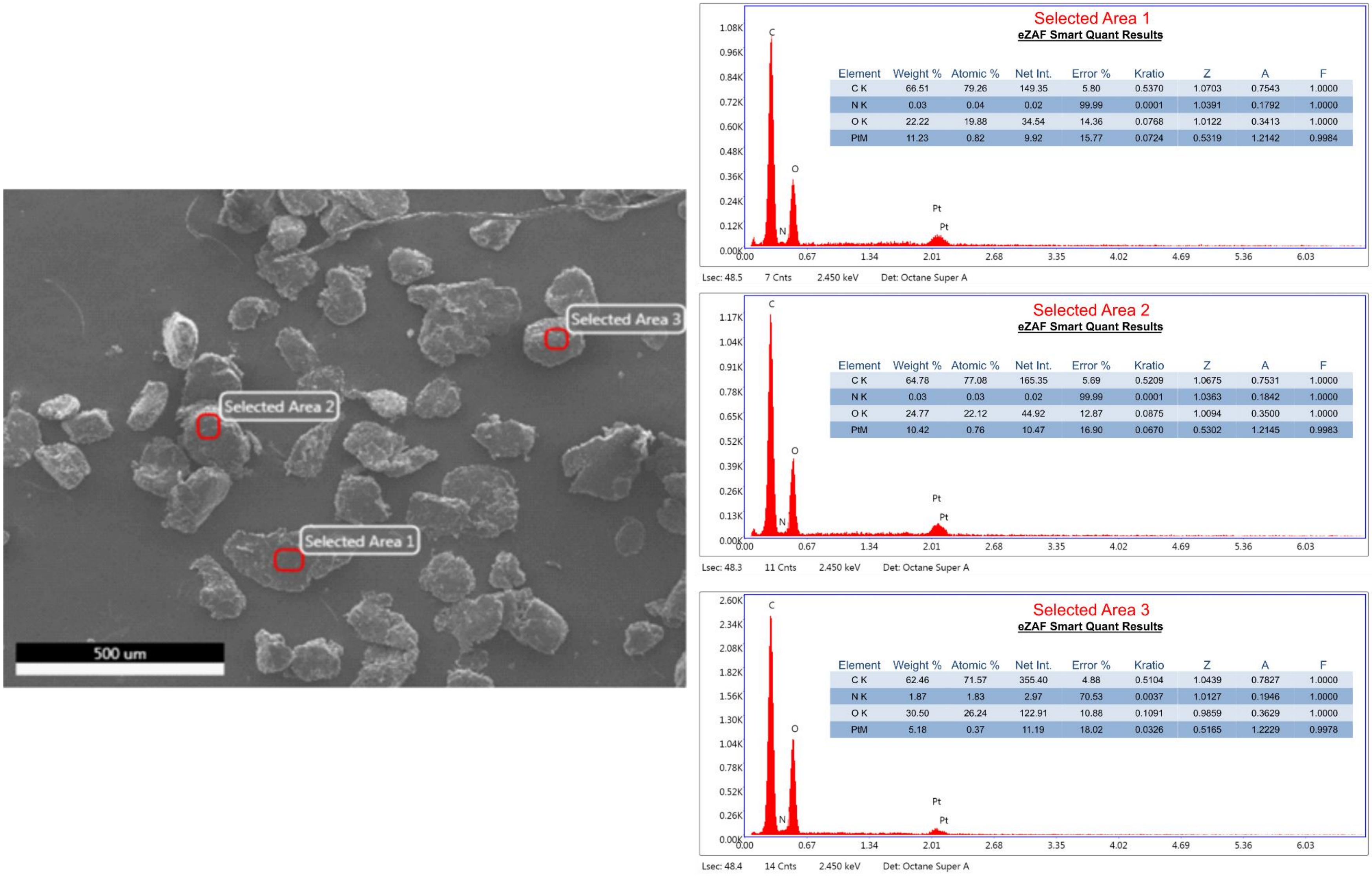

**Fig. S9.**

SEM image of the washed PET material after the 1$^{st}$ round of hydrolysis (see ***Multi-round RAging experiments***), and the corresponding EDX spectra indicating that the remaining PET surfaces have a varying amount of nitrogen present (from the protein).

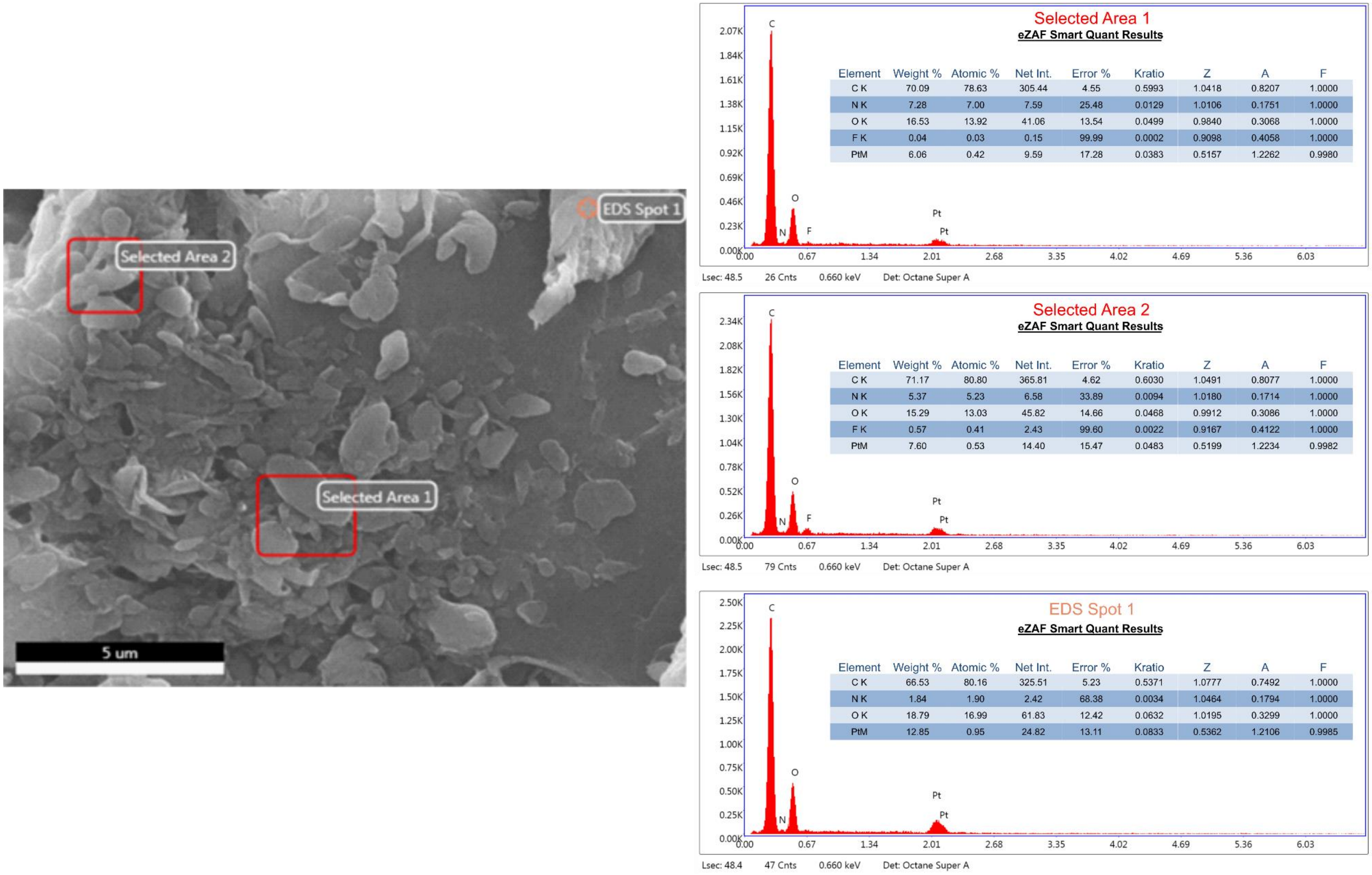

Selected Area 1 — eZAF Smart Quant Results

| Element | Weight % | Atomic % | Net Int. | Error % | Kratio | Z | A | F |
|---|---|---|---|---|---|---|---|---|
| C K | 70.09 | 78.63 | 305.44 | 4.55 | 0.5993 | 1.0418 | 0.8207 | 1.0000 |
| N K | 7.28 | 7.00 | 7.59 | 25.48 | 0.0129 | 1.0106 | 0.1751 | 1.0000 |
| O K | 16.53 | 13.92 | 41.06 | 13.54 | 0.0499 | 0.9840 | 0.3068 | 1.0000 |
| F K | 0.04 | 0.03 | 0.15 | 99.99 | 0.0002 | 0.9098 | 0.4058 | 1.0000 |
| PtM | 6.06 | 0.42 | 9.59 | 17.28 | 0.0383 | 0.5157 | 1.2262 | 0.9980 |

Selected Area 2 — eZAF Smart Quant Results

| Element | Weight % | Atomic % | Net Int. | Error % | Kratio | Z | A | F |
|---|---|---|---|---|---|---|---|---|
| C K | 71.17 | 80.80 | 365.81 | 4.62 | 0.6030 | 1.0491 | 0.8077 | 1.0000 |
| N K | 5.37 | 5.23 | 6.58 | 33.89 | 0.0094 | 1.0180 | 0.1714 | 1.0000 |
| O K | 15.29 | 13.03 | 45.82 | 14.66 | 0.0468 | 0.9912 | 0.3086 | 1.0000 |
| F K | 0.57 | 0.41 | 2.43 | 99.60 | 0.0022 | 0.9167 | 0.4122 | 1.0000 |
| PtM | 7.60 | 0.53 | 14.40 | 15.47 | 0.0483 | 0.5199 | 1.2234 | 0.9982 |

EDS Spot 1 — eZAF Smart Quant Results

| Element | Weight % | Atomic % | Net Int. | Error % | Kratio | Z | A | F |
|---|---|---|---|---|---|---|---|---|
| C K | 66.53 | 80.16 | 325.51 | 5.23 | 0.5371 | 1.0777 | 0.7492 | 1.0000 |
| N K | 1.84 | 1.90 | 2.42 | 68.38 | 0.0034 | 1.0464 | 0.1794 | 1.0000 |
| O K | 18.79 | 16.99 | 61.83 | 12.42 | 0.0632 | 1.0195 | 0.3299 | 1.0000 |
| PtM | 12.85 | 0.95 | 24.82 | 13.11 | 0.0833 | 0.5362 | 1.2106 | 0.9985 |

**Fig. S10.**

SEM image of the washed PET material after 7 rounds of hydrolysis (see ***Multi-round RAging experiments***), and the corresponding EDX spectra indicating that the remaining PET surfaces have a varying amount of nitrogen present (from the protein).

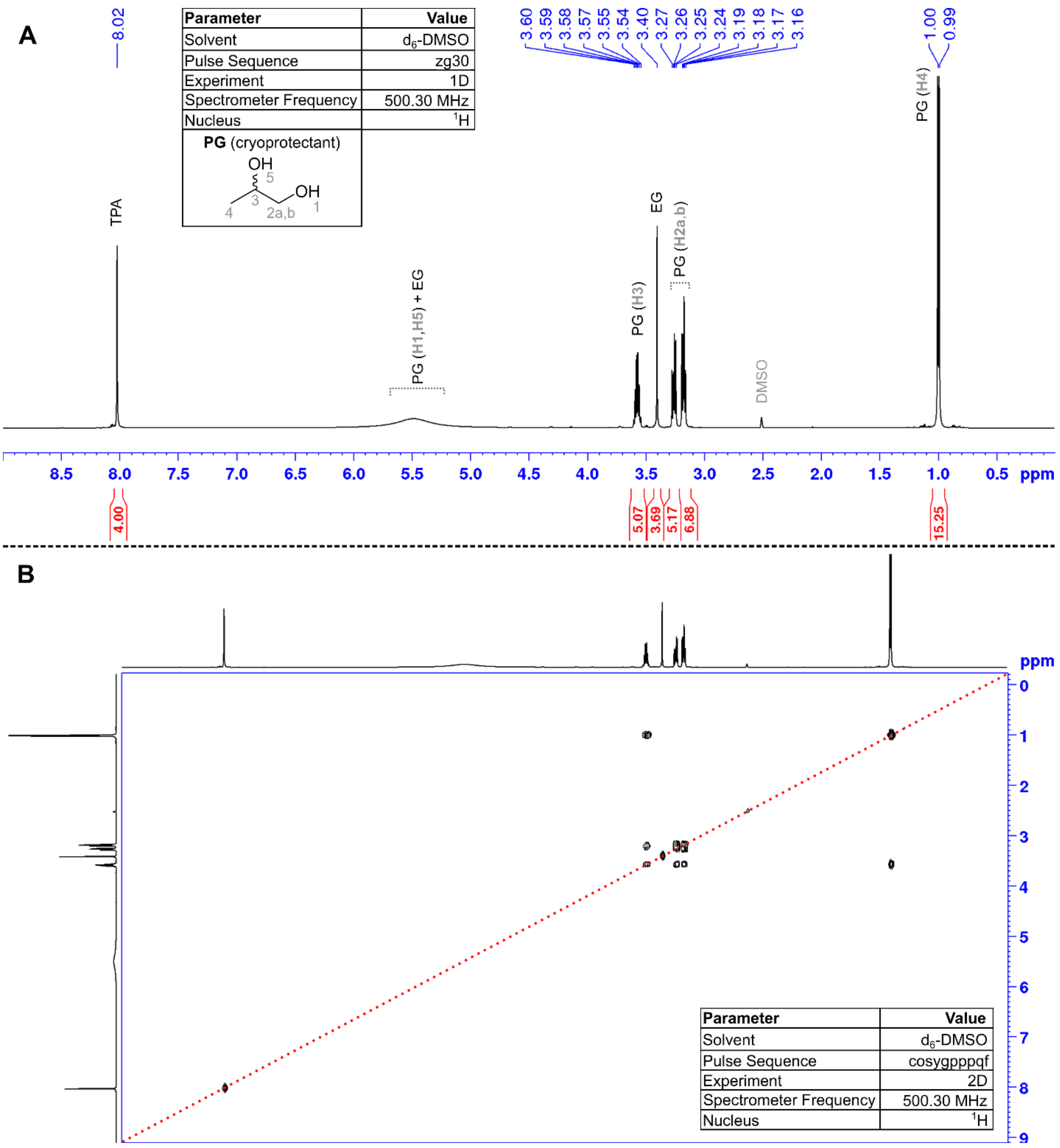

**Fig. S11.**

The $^{1}H$ NMR spectra of the product extracted from the reaction mixture using extraction method A (See SI Methods: ***TPA isolation***). The 1D (A) and 2D (B) spectra show that the product contains, in addition to TPA and EG, the cryoprotectant 1,2-propylene glycol originating from the commercial enzyme preparation. The very broad signal for the acidic protons (2H) of TPA at 13.29 ppm is the only signal outside the range shown.

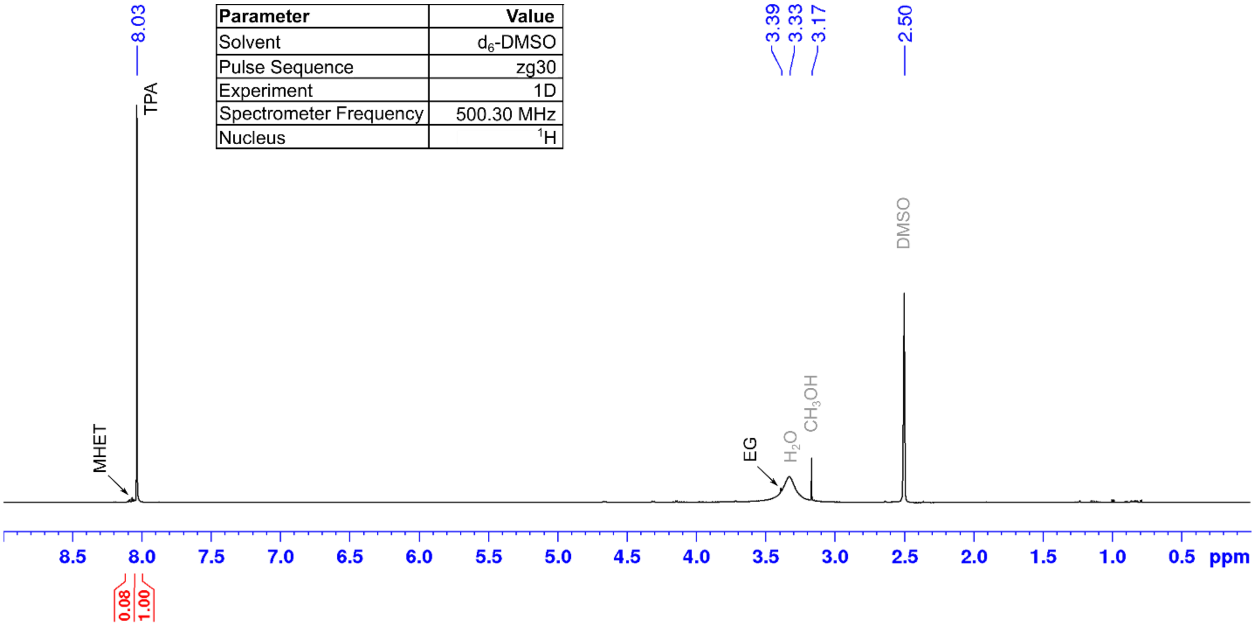

| Parameter | Value |
|---|---|
| Solvent | $d_6$-DMSO |
| Pulse Sequence | zg30 |
| Experiment | 1D |
| Spectrometer Frequency | 500.30 MHz |
| Nucleus | $^1H$ |

**Fig. S12.**

The $^1$H NMR spectra of the product extracted from the reaction mixture using method B (See SI Methods: ***TPA isolation***). The very broad signal for the acidic protons (2H) of TPA at 13.29 ppm is the only signal outside the range shown.

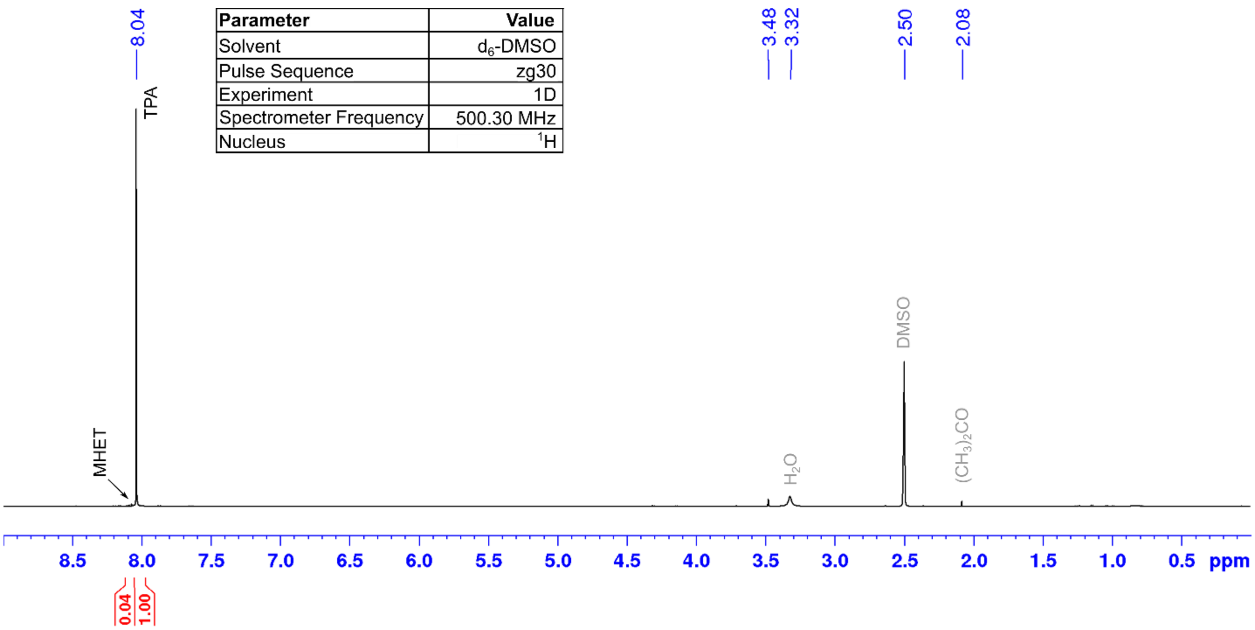

**Fig. S13.**

The $^{1}H$ NMR spectra of the product extracted from the reaction mixture using method C (See SI Methods: ***TPA isolation***). The broad signal for the acidic protons (2H) of TPA at 13.29 ppm is the only signal outside the range shown. The small acetone peak at 2.08 ppm is likely from insufficient drying of the used NMR tube. The signal at 3.48 ppm does not correspond to ethylene glycol nor to any expected solvent impurity.

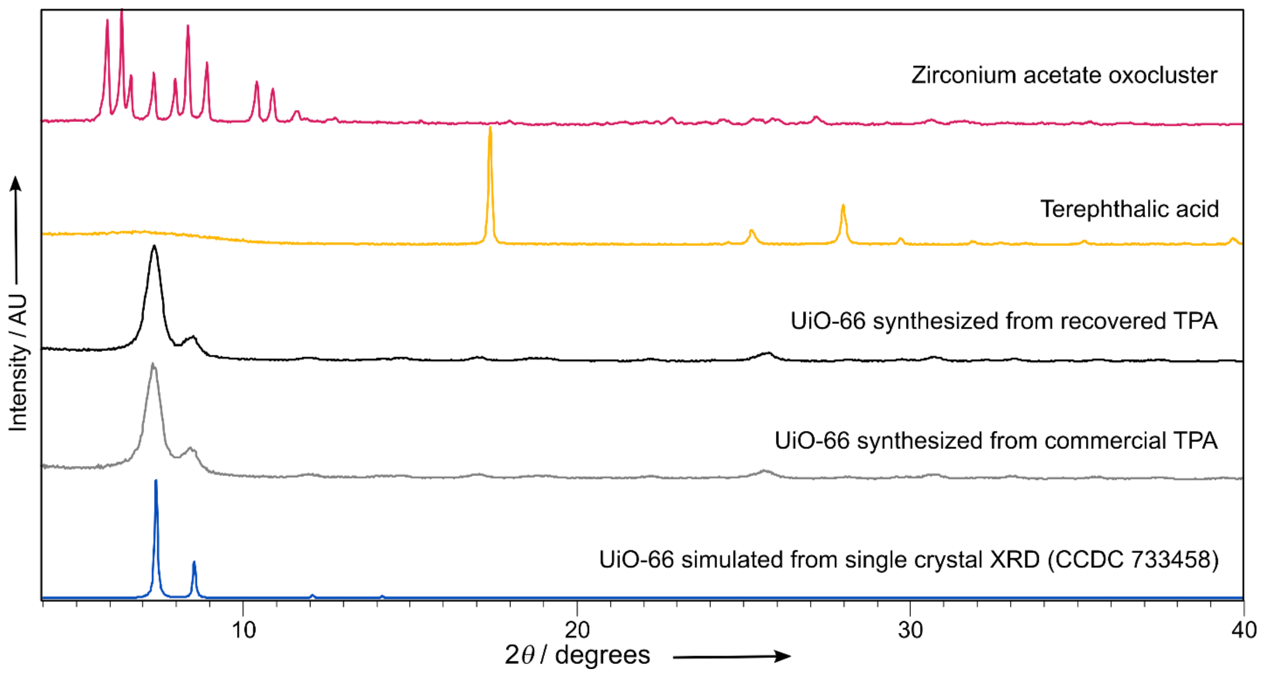

**Fig. S14.**

PXRD patterns of the starting materials: the zirconium acetate oxocluster $Zr_{12}O_8(OH)_8(CH_3COO)_{24}$ (in magenta) and TPA (in yellow); the obtained UiO-66 from the recovered TPA (in black) and commercially obtained TPA (in gray) and the simulated powder pattern from the crystal structure of UiO-66 (CCDC 733485) from ref (*5*).

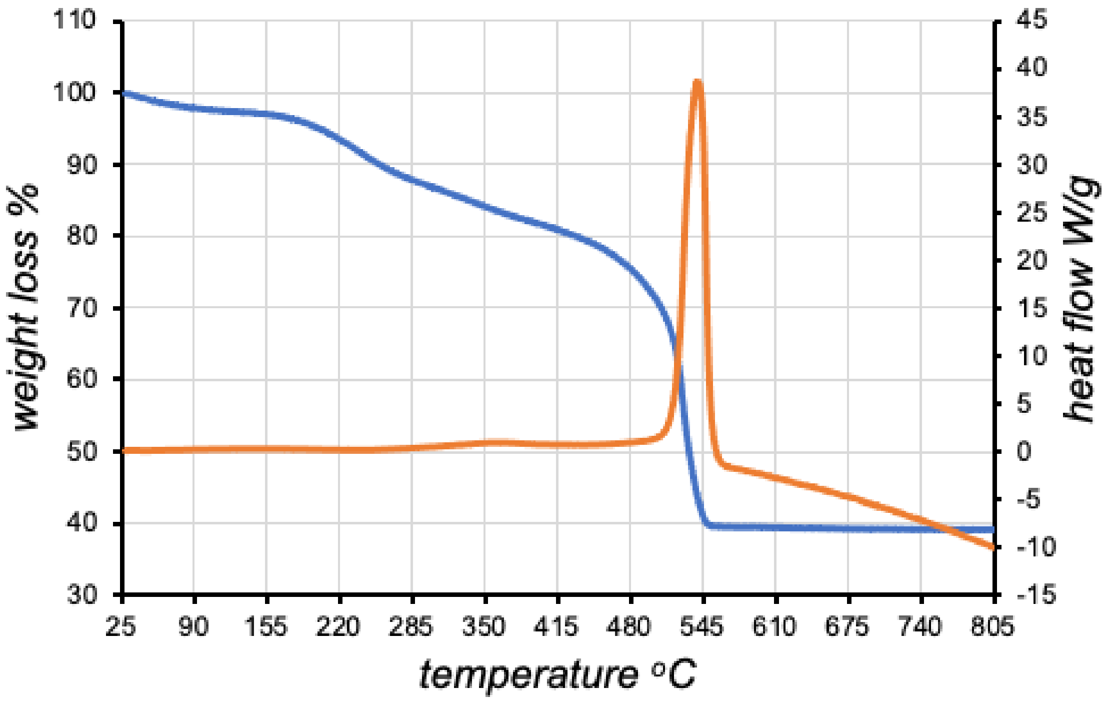

**Fig. S15.**

TG and DSC analysis of the UiO-66 synthesized from the **TPA recycled from PET** by enzymatic depolymerization in damp-solid reaction mixtures. The synthesized UiO-66 was washed thoroughly with methanol (see SI ***Synthesis of the UiO-66 metal-organic framework (MOF)***) and dried at 80°C in vacuum for 36 h prior to TG/DSC measurement. The total sample weight was 4.498 mg. The residual solvent loss was observed between 50°C and 120°C, followed by a gradual mass loss between 140 and 400°C corresponding to the dehydration of the $Zr_6O_4(OH)_4$ nodes to $Zr_6O_6$. The sharp mass loss (2.101 mg) between 320–650°C represents the decomposition of the organic linker TPA (exp. 46.7%, calc. for 6 linkers is 60.2%), which corresponds to the existence of 4.65 mg of TPA in the UiO-66 structure. The residue of $ZrO_2$ is 39.3%.

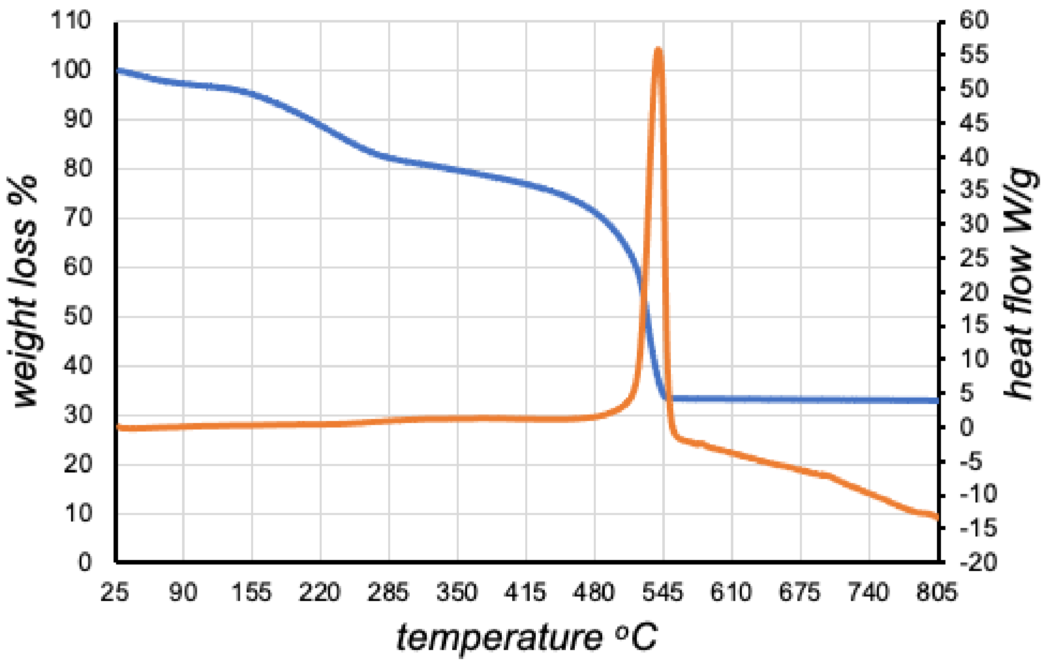

**Fig. S16.**

TGA and DSC analysis of the UiO-66 synthesized from **commercially obtained TPA**. The synthesized UiO-66 was washed thoroughly with methanol (see SI ***Synthesis of the UiO-66 metal-organic framework (MOF)***) and dried at 80°C in vacuum for 36 h prior to TG/DSC measurement. The total sample weight was 4.39 mg. The solvent loss was observed between 50°C and 120°C, followed by a gradual mass loss between 140 and 400°C corresponding to the dehydration of the $Zr_6O_4(OH)_4$ nodes to $Zr_6O_6$. The sharp mass loss (2.089 mg) between 320–650°C represents the decomposition of the organic linker TPA (exp. 47.6%, calc. for 6 linkers is 60.2%), which corresponds to the existence of 4.72 mg of TPA in the UiO-66 structure. The residue of $ZrO_2$ is 33.2%.

### Supplementary Tables

#### Table S1.

Conditions varied and results for the reactions shown on Scheme S1.

| Entry | PET (mg) | Buffer added | $\eta$ (µL mg$^{-1}$) | Enz. loading | Milling duration (min) | Aging temp. (°C) | Aging duration (days) | Yield of **TPA** (%) |
|---|---|---|---|---|---|---|---|---|
| 1 | 300 | 0.1 M Na-PB pH 7.3 [a] | 1.5 | 0.6 wt% | **0** | 55 | 7 | 19 ± 1 |
| 2 | 300 | 0.1 M Na-PB pH 7.3 | 1.5 | 0.6 wt% | **1** | 55 | 7 | 24 ± 1 |
| 3 | 300 | 0.1 M Na-PB pH 7.3 | 1.5 | 0.6 wt% | **2** | 55 | 7 | 25 ± 6 |
| 4 | 300 | 0.1 M Na-PB pH 7.3 | 1.5 | 0.6 wt% | **5** | 55 | 7 | 20 ± 1 |
| 5 | 300 | 0.1 M Na-PB pH 7.3 | 1.5 | 0.6 wt% | **10** | 55 | 7 | 17.6 ± 0.6 |
| 6 | 300 | 0.1 M Na-PB pH 7.3 | 1.5 | 0.6 wt% | **30** | 55 | 7 | 16.3 ± 0.7 |
| 7 | 300 | 0.1 M Na-PB pH 7.3 | 1.5 | 0.6 wt% | 5 | **35** | 7 | 8.0 ± 0.4 |
| 8 | 300 | 0.1 M Na-PB pH 7.3 | 1.5 | 0.6 wt% | 5 | **40** | 7 | 14.8 ± 0.8 |
| 9 | 300 | 0.1 M Na-PB pH 7.3 | 1.5 | 0.6 wt% | 5 | **45** | 7 | 23 ± 3 |
| 10 | 300 | 0.1 M Na-PB pH 7.3 | 1.5 | 0.6 wt% | 5 | **50** | 7 | 24 ± 3 |
| 11 | 300 | 0.1 M Na-PB pH 7.3 | 1.5 | 0.6 wt% | 5 | **55** | 7 | 22.6 ± 0.9 |
| 12 | 300 | 0.1 M Na-PB pH 7.3 | 1.5 | 0.6 wt% | 5 | **60** | 7 | 3.0 ± 0.4 |
| 13 | 300 | 0.1 M Na-PB pH 7.3 | 1.5 | 0.6 wt% | 5 | **65** | 7 | 1.7 ± 0.2 |
| 14[b] | 300 | 0.1 M Na-PB pH 7.3 | 1.5 | **0.6 wt% boiled** | 5 | 55 | 7 | 1.78 ± 0.06 [c] |
| 15 | 300 | 0.1 M Na-PB **pH 6** | 1.5 | 0.6 wt% | 5 | 55 | 7 | 17 ± 1 |
| 16 | 300 | 0.1 M Bicine **pH 8** | 1.5 | 0.6 wt% | 5 | 55 | 7 | 17.9 ± 0.2 |
| 17 | 300 | 0.1 M Bicine **pH 9** | 1.5 | 0.6 wt% | 5 | 55 | 7 | 20 ± 1 |
| 18 | 300 | **MilliQ** | 1.5 | 0.6 wt% | 5 | 55 | 7 | 14 ± 1 |
| 19 | 300 | **0.5 M Bicine pH 9** | 1.5 | 0.6 wt% | 5 | 55 | 7 | 24.6 ± 0.6 [c] |
| 20 | 300 | **1 M Tris-HCl pH 8.2** | 1.5 | 0.6 wt% | 5 | 55 | 7 | 21.0 ± 0.4 |
| 21 | **600** | 0.1 M Na-PB pH 7.3 | 1.5 | 0.6 wt% | 5 | 55 | 7 | 19 ± 2 |
| 22 | 300 | 0.1 M Na-PB pH 7.3 | 1.5 | **0.6 wt% pre-aged[e]** | 5 | 55 | 7 | 18 [d] |
| 23 | 300 | 0.5 M Bicine pH 9 | 1.5 | **0.6 wt% pre-aged[e]** | 5 | 55 | 7 | 24 [d] |
| **Controls** | | | | | | | | |
| 24 (in-solution) | 8–12 | 0.1 M Na-PB pH 7.3 | **50** | 0.6 wt% | – | 55 | 7 | 10 ± 1 [f] |
| 25 (blank) | 300 | 0.1 M Na-PB pH 7.3 | 1.5 | **0.0 wt%** | **30** | 55 | 7 | 0 [d] |

[a] 0.1 M sodium phosphate buffer, pH 7.3. [b] The 300 µL of enzyme solution used was boiled for 30 minutes in a water bath prior to the reaction. [c] Reaction carried out in duplicate. [d] Reaction carried out in monoplicate. [e] The enzyme mixed with buffer was kept at 55°C for 3 days before the reaction was started. [f] The reaction produces also 3.6 ± 0.7 % yield of MHET, giving the solution reaction 2.7-fold selectivity of TPA over MHET.

**Table S2.**

Conditions varied and results for the reactions shown on Scheme S2, using the lyophilized enzyme.

| Entry | PET (mg) | Buffer added | $\eta$ (µL mg$^{-1}$) | Enz. loading[a] | Milling duration (min) | Aging temp. (°C) | Aging duration (days) | Yield of **TPA** (%) |
|---|---|---|---|---|---|---|---|---|
| 1 | 300 | **1 M Tris-HCl pH 8.2** | 1.5 | 0.6 wt% (lyo) | 5 | 55 | 7 | 18 ± 3 [b] |
| 2 | 300 | MilliQ | **0.5** | 0.6 wt% (lyo) | 5 | 55 | 7 | 13.2 ± 0.4 |
| 3 | 300 | MilliQ | **1.0** | 0.6 wt% (lyo) | 5 | 55 | 7 | 15 ± 1 |
| 4 | 300 | MilliQ | **1.5** | 0.6 wt% (lyo) | 5 | 55 | 7 | 14.5 ± 0.6 |
| 5 | 300 | MilliQ | **2.0** | 0.6 wt% (lyo) | 5 | 55 | 7 | 15.0 ± 0.6 |
| 6 | 300 | 0.1 M Na-PB pH 7.3 + MilliQ[c] | **0.5** | 0.6 wt% (lyo) | 5 | 55 | 7 | 15.4 ± 0.3 |
| 7 | 300 | 0.1 M Na-PB pH 7.3 + MilliQ[d] | **1.0** | 0.6 wt% (lyo) | 5 | 55 | 7 | 14 ± 1 |
| 8 | 300 | 0.1 M Na-PB pH 7.3 + MilliQ[e] | **1.5** | 0.6 wt% (lyo) | 5 | 55 | 7 | 18 ± 1 |
| 9 | 300 | 0.1 M Na-PB pH 7.3 + MilliQ[f] | **2.0** | 0.6 wt% (lyo) | 5 | 55 | 7 | 14.5 ± 0.8 |

[a] 16–17 mg of the lyophilized powder was used per reaction, which contains 12 wt% HiC, 11 wt% Tris-HCl pH 7.3 buffer and 89 wt% NaCl. [b] Reaction carried out in duplicate. [c] 75 µL of 0.1 M sodium phosphate buffer and 75 µL of water. [d] 150 µL of 0.1 M sodium phosphate buffer and 150 µL of water. [e] 150 µL of 0.1 M sodium phosphate buffer and 300 µL of water. [f] 150 µL of 0.1 M sodium phosphate buffer and 450 µL of water.

**Table S3.**

Conditions varied and results for reactions shown on Scheme S1 (other PET sources)

| Entry | PET (mg) | PET source | Crystallinity (%) | Buffer added | $\eta$ (µL mg$^{-1}$) | Enz. loading | Milling duration (min) | Aging temp. (°C) | Aging duration (days) | Yield of **TPA** (%) |
|---|---|---|---|---|---|---|---|---|---|---|
| 1 | 300 | **Bottle (pre-milled)** | 31% | 0.1 M Na-PB pH 7.3 [a] | 1.5 | 0.6 wt% | 30[b] | 55 | 7 | 16 ± 2 |
| 2 | 300 | **Bottle squares** | 30%[c] | 0.1 M Na-PB pH 7.3 | 1.5 | 0.6 wt% | 30 | 55 | 7 | 15 ± 4 |
| 3 | 300 | **Bottle + cap (pre-milled)** | 31% [d] | 0.1 M Na-PB pH 7.3 | 1.5 | 0.6 wt% | 30 | 55 | 7 | 19 [e] |
| 4 | 300 | **Container, 80% rPET (pre-milled)** | 34% | 0.1 M Na-PB pH 7.3 | 1.5 | 0.6 wt% | 5 | 55 | 7 | 15.± 1 |
| 5 | 300 | **Thin film (pre-milled)** | 32% | 0.1 M Na-PB pH 7.3 | 1.5 | 0.6 wt% | 5 | 55 | 7 | 17 ± 1 |
| 6 | 300 | **Textile, 100% polyester** | 46% | 0.1 M Na-PB pH 7.3 | 1.5 | 0.6 wt% | 5 | 55 | 7 | 6.3 ± 0.1 [f] |

[a] 0.1 M sodium phosphate buffer, pH 7.3. [b] Reaction carried out in a 15 mL Teflon jar. [c] the higher error arises in part from the fact that the the bottle plastic varies in thickness. Pre-milled powder (that gives a lower error) is a better average of all the bottle parts. [d] Assumed based on the same PET pre-milling conditions as entry 1, the crystallinity of mixed polymer was not calculated based on the DSC curve (Figure S3). [e] Reaction carried out in monoplicate. [f] Reaction carried out in duplicate.

**Table S4.**

Effect from non-participating polymers on the reaction shown on Scheme S1.

| Entry | PET (mg) | Non-participating polymer | Buffer added | $\eta$ (µL mg$^{-1}$) | Enz loading | Milling duration (min) | Aging temp. (°C) | Aging duration (days) | Yield of **TPA** (%) |
|---|---|---|---|---|---|---|---|---|---|
| 1 | 100 | **200 mg Polystyrene** | 0.1 M Na-PB pH 7.3 [a] | 1.5 | 0.6 wt% | **5** | 55 | 7 | 19[c] |
| 2 | 100 | **200 mg MCC[b]** | 0.1 M Na-PB pH 7.3 | 1.5 | 0.6 wt% | **5** | 55 | 7 | 16[c] |

[a] 0.1 M sodium phosphate buffer, pH 7.3. [b] Micro-crystalline cellulose. [c] Reaction carried out in monoplicate.

**Table S5.**

Conditions varied and results of testing the scope of the reaction shown on Scheme S1.

| Entry | Plastic (mg) | Plastic type | Buffer added | $\eta$ (µL mg$^{-1}$) | Enz. loading | Milling duration (min) | Aging temp. (°C) | Aging duration (days) | Yield of hydrolysis (%) |
|---|---|---|---|---|---|---|---|---|---|
| 1 | 300 | **Polybutylene terephthalate (PBT)** | 0.1 M Na-PB pH 7.3[a] | 1.5 | 0.6 wt% | 30[a] | 55 | 7 | 0.9 ± 0.4[b] |
| 2 | 300 | **Polycarbonate (PC)** | 0.1 M Na-PB pH 7.3 | 1.5 | 0.6 wt% | 30 | 55 | 7 | 3.81 ± 0.03[c] |
| 3 | 300 | **Polycaprolactone (PCL)** | 0.1 M Na-PB pH 7.3 | 1.5 | 0.6 wt% | 30 | 55 | 7 | TBD[d] |
| **Controls** | | | | | | | | | |
| 4 | 8–12 | **PBT** | 0.1 M Na-PB pH 7.3 | 50 | 0.6 wt% | 30[a] | 55 | 7 | 0.70 ± 0.03[b] |
| 5 | 8–12 | **PC** | 0.1 M Na-PB pH 7.3 | 50 | 0.6 wt% | 30 | 55 | 7 | 2.2 ± 0.1[c] |
| 6 | 8–12 | **PCL** | 0.1 M Na-PB pH 7.3 | 50 | 0.6 wt% | 30 | 55 | 7 | TBD[d] |

[a] 0.1 M sodium phosphate buffer, pH 7.3. [b] Calculated from theoretical amount of TPA monomer produced. [c] Calculated from theoretical amount of BPA monomer produced. [d] Calculated from theoretical amount of 6-hydroxycaproic acid monomer produced.

**Table S6.**

Conditions varied and results for the RAging reactions shown on Scheme S3.

| Entry | PET (mg) | Buffer added | $\eta$ (µL mg$^{-1}$) | Enz loading | RAging regime (milled + aged) | Number of cycles | Yield of **TPA** (%) |
|---|---|---|---|---|---|---|---|
| 1 | 300 | 0.1 M Na-PB pH 7.3 [a] | 1.5 | 0.6 wt% | 5 min + 1 day 55°C | 3 | 21 ± 2 |
| | | | | | | 7 | 22 ± 3 |
| 2 | **900** | 0.1 M Na-PB pH 7.3 | 1.5 | 0.6 wt% | 5 min + 1 day 55°C | 3 | 21.5 ± 0.5 |
| | | | | | | 7 | 26 ± 2 |
| 3 | 300 | 0.1 M Na-PB pH 7.3 | 1.5 | **2.6 wt%** | 5 min + 1 day 55°C | 3 | 18 ± 2 |
| | | | | | | 7 | 25.4 ± 0.9 |
| **Control** | | | | | | | |
| blank | 300 | 0.1 M Na-PB pH 7.3 | 1.5 | **0 wt%** | 5 min + 1 day 55°C | 7 | 0[b][c] |

[a] 0.1 M sodium phosphate buffer, pH 7.3. [b] Reaction was carried out in monoplicate. [c] The reaction mixture did not contain any detectable amount of TPA after 7 days of RAging, based on HPLC analysis.

**Table S7.**

Conditions varied and results for product inhibition studies (See SI ***Method for the product inhibition studies***).

| Entry | PET (mg) | Additive, amount | Buffer added | $\eta$ (µL mg$^{-1}$) | Enz. loading | Milling duration (min) | Aging temp. (°C) | Aging duration (days) | Yield of **TPA** (%) |
|---|---|---|---|---|---|---|---|---|---|
| 1 | 250 | **TPA, 50 mg** | 0.1 M Na-PB pH 7.3 [a] | 1.5 | 0.6 wt% | 5 | 55 | 7 | 19 ± 2[b] |
| 2 | 250 | **TPA, 50 mg + EG 20 µL** | 0.1 M Na-PB pH 7.3 | 1.5 | 0.6 wt% | 5[c] | 55 | 7 | 18.5 ± 0.5[b] |
| 3 | 300 | **EG, 150 µL** | MilliQ | 2.0 | 0.6 wt% | 5[d] | 55 | 7 | 22[e] |
| 4 | 300 | **EG, 150 µL** | MilliQ | 2.0 | **0 wt%** | 5[d] | 55 | 7 | 0[e] |

[a] 0.1 M sodium phosphate buffer, pH 7.3. [b] The amount of TPA present from the start of the reaction is subtracted from the total amount of TPA contained in the reaction mixture after 7 days of aging, therefore the conversion reported here only accounts for the TPA generated during the reaction. [c] PET, TPA and EG were pre-milled together for 1 minute before other reaction components were added, to ensure better mixing of the liquid EG and solids. [d] PET and EG were pre-milled together for 10 minutes before other reaction components were added, to ensure better mixing of the liquid EG and solid PET. [e] Reaction was carried out as monoplicate.

### Table S8.

Results of the BHET enzyme activity assays: Novozym® 51032 as purchased (entries 1, 2), extracted into the reaction mixture washings (entries 3–7) and remaining on solids (entries 8, 9).

| Entry | Extr. from | Extr. when | Assay buffer | Remaining PET | Assay duration (min) | Assay temp. (°C) | BHET consumed (%) | Enzyme activity (U)[a] | Activity remaining |
|---|---|---|---|---|---|---|---|---|---|
| 1 | - | - | 0.1 M Na-PB pH 7.3 [b] | - | 5 | 55 | 98.2 ± 0.5 | 0.0196 ± 0.0001 | **100%** |
| 2 [c] | - | - | 0.1 M Na-PB pH 7.3 | - | 5 | 55 | 97.5 ± 0.7 | 0.0195 ± 0.0001 | **99%** |
| 3 | PET | Milled | 0.1 M Na-PB pH 7.3 | - | 5 | 55 | 95 ± 2 | 0.0189 ± 0.0004 | **97%** |
| 4 | PET | Day 3 | 0.1 M Na-PB pH 7.3 | - | 10 | 55 | 0 | 0 | **0%** |
| 5 | PET | Day 7 | 0.1 M Na-PB pH 7.3 | - | 10 | 55 | 0 | 0 | **0%** |
| 6 | PBT | Day 3 | 0.1 M Na-PB pH 7.3 | - | 10 | 55 | 99 ± 1 | 0.0099 ± 0.0001 | **51%** |
| 7 | PBT | Day 7 | 0.1 M Na-PB pH 7.3 | - | 10 | 55 | 97.5 ± 0.6 | 0.0098 ± 0.0001 | **50%** |
| 8 | - | - | 0.1 M Na-PB pH 7.3 | 10 mg[d] | 120 | 55 | 78 ± 5 | 0.0007 | **4%** |
| 9 | - | - | 0.1 M Na-PB pH 7.3 | 10 mg[e] | 120 | 55 | 94 ± 4 | 0.0008 | **4%** |

[a] The activity unit U is based on the amount of BHET consumed during the assay, U = μmol(BHET) $min^{-1}$. [b] 0.1 M sodium phosphate buffer, pH 7.3. [c] Novozym® 51032 after storage for 6 months [d] The reaction mixture was washed with methanol to recover the remaining PET. [e] The reaction mixture was washed with Tris-HCl buffer (pH 8.2) and water to recover the remaining PET. The details of the solid's assays are described in the section: ***BHET enzyme activity assay on the HiC stuck on remaining hydrolyzed PET solids.***

**Table S9.**

Space-time yield and TPA production from the current and previous work using enzymes to depolymerize PET

| Enzyme | Organism | PET source | PET crystallinity | Volume (mL) | PET loading (mg mL$^{-1}$) | Enz. loading (mg mL$^{-1}$) | Reaction temp. (˚C) | Time (h) | TPA yield (mg mL$^{-1}$) | Space-time yield ($g_{TPA}$ L$^{-1}$ h$^{-1}$) | TPA per gram of enz. ($g_{TPA}$ $g_{enz.}^{-1}$ h$^{-1}$) | Reference |
|---|---|---|---|---|---|---|---|---|---|---|---|---|
| Thc_Cut1 | *Thermobifida cellulosilytica* | PET film | 37% | 0.4 | not reported | 0.2 | 50 | 120 | $5.61 \times 10^{-5}$ | 0.00 | 0.00 | (*6*) |
| Thc_Cut2 | *Thermobifida cellulosilytica* | PET film | 37% | 0.4 | not reported | 0.2 | 50 | 120 | $5.61 \times 10^{-6}$ | 0.00 | 0.00 | (*6*) |
| Thf42_Cut1 | *Thermobifida fusca* | PET film | 37% | 0.4 | not reported | 0.2 | 50 | 120 | $4.60 \times 10^{-5}$ | 0.00 | 0.00 | (*6*) |
| Tfu_0883 | *Thermobifida fusca* | PET fabric | not reported | 100 | 4 | 0.05 | 60 | 2 | 1.20[a] | 0.60 | 12.00 | (*7*) |
| Tfu_0883 (I218A) | *Thermobifida fusca* | PET fabric | not reported | 100 | 4 | 0.05 | 60 | 2 | 2.54[a] | 1.27 | 25.40 | (*7*) |
| Tfu_0883 (Q132A/T101A) | *Thermobifida fusca* | PET fabric | not reported | 100 | 4 | 0.05 | 60 | 2 | 2.81[a] | 1.41 | 28.10 | (*7*) |
| Thc_Cut1 | *Thermobifida cellulosilytica* | PET film | not reported | 1 | not reported | 0.2 | 50 | 48 | 0.04 | 0.00 | 0.00 | (*8*) |
| Thc_Cut2 | *Thermobifida cellulosilytica* | PET film | not reported | 1 | not reported | 0.2 | 50 | 48 | 0.01 | 0.00 | 0.00 | (*8*) |
| Thc_Cut2 (R29N/A30V) | *Thermobifida cellulosilytica* | PET film | not reported | 1 | not reported | 0.2 | 50 | 48 | 0.07 | 0.00 | 0.01 | (*8*) |
| Thc_Cut2 (R19S/R29N/ A30V) | *Thermobifida cellulosilytica* | PET film | not reported | 1 | not reported | 0.2 | 50 | 48 | 0.06 | 0.00 | 0.01 | (*8*) |
| Cut190 | *Saccharomonospora viridis AHK190* | PET film | not reported | 1 | 25 | 0.06 | 48 | 72 | 0.0019 | 0.00 | 0.00 | (*9*) |
| Cut190 | *Saccharomonospora viridis AHK190* | PET packaging | not reported | 1 | 25 | 0.06 | 48 | 72 | 0.0045 | 0.00 | 0.00 | (*9*) |
| TfCut2 | *Thermobifida fusca* | PET nanoparticles | 9.8% | 1 | not reported | 0.05 | 60 | 24 | $9.97 \times 10^{-5}$ | 0.00 | 0.00 | (*10*) |
| TfCa+TfCut2 | *Thermobifida fusca KW3* | PET film | not reported | 40 | 162.5 | 0.04 | 60 | 24 | 0.58 | 0.02 | 0.60 | (*11*) |
| TfCa+LCC | Leaf-branch compost meta-genomic study | PET film | not reported | 40 | 162.5 | 0.04 | 60 | 24 | 1.29 | 0.05 | 1.34 | (*11*) |
| TfCut2 (G62A) | *Thermobifida fusca KW3* | PET film | not reported | 1.8 | 25 | 0.03 | 65 | 50 | 0.012 | 0.00 | 0.01 | (*12*) |
| PETase | *Ideonella sakaiensis 201-F6* | PET film | 1.9% | 0.3 | not reported | 0.001 | 30 | 18 | 0.10 | 0.01 | 5.56 | (*13*, *14*) |

| | | | | | | | | | | | | |
|---|---|---|---|---|---|---|---|---|---|---|---|---|
| PETase | *Ideonella sakaiensis 201-F6* | high crystallinity PET film | not reported | 0.3 | not reported | 0.001 | 30 | 18 | $6.65 \times 10^{-4}$ | 0.00 | 0.04 | (*13*, *14*) |
| CALB/HiC | *Candida antarctica/ Humicola insolens* | PET water bottle | 36.5% | 5 | 40 | 1.6 | 50 | 336 | 0.37 | 0.00 | 0.00 | (*15*) |
| CALB/HiC | *Candida antarctica/ Humicola insolens* | PET film | 4.9% | 10 | 20 | 0.4 | 60˚C (14d), then 37˚C (1d) | 360 | 9.97 | 0.03 | 0.07 | (*16*) |
| CALB/HiC | *Candida antarctica/ Humicola insolens* | PET flakes | 41.1% | 10 | 20 | 0.4 | 60˚C (14d), then 37˚C (1d) | 360 | 2.26 | 0.01 | 0.02 | (*16*) |
| HiC | *Humicola insolens* | Treated PET fibers | not reported | 2 | 5 | 1 | 50 | 6 | 1.08 | 0.18 | 0.18 | (*17*) |
| PETase (W159H/S238F) | *Ideonella sakaiensis 201-F6* | PET film | 14.8% | 0.5 | not reported | 0.001 | 30 | 96 | 0.068 | 0.00 | 0.71 | (*18*) |
| PETase (Y58A) | *Ideonella sakaiensis 201-F6* | PET bottle | not reported | 0.7 | not reported | 0.001 | 30 | 20 | $3.49 \times 10^{-6}$ | 0.00 | 0.00 | (*19*) |
| PETase (I179F) | *Ideonella sakaiensis 201-F6* | PET film | not reported | 1.5 | not reported | 0.003 | 30 | 48 | 1.06 | 0.02 | 7.36 | (*20*) |
| HiC | *Humicola insolens* | Post-consumer PET | 41.1% | 10 | 80.8 | 5.24 | 62.6 | 168 | 16.8 | 0.10 | 0.02 | (*21*) |
| LCC (Y127G/D238C/ F243I/S283C) | Leaf-branch compost meta-genomic study | Thermomechani cally treated Post-consumer PET | 14.6% [b] | 80 | 250 | 0.75 | 72 | 12.5 | 208.6 | 16.7 | 22.25 | (*22*) |
| **HiC** | ***Humicola insolens*** | **PET powder** | **36.2%** | **0.45** | **666.7** | **4.3** | **55** | **72** | **115.6** | **1.61** | **0.37** | **This work**[c] |
| **HiC** | ***Humicola insolens*** | **PET powder** | **36.2%** | **1.35** | **666.7** | **30.1**[d] | **55** | **504** | **281.4** | **0.56** | **0.02** | **This work**[e] |
| **HiC** | ***Humicola insolens*** | **PET powder** | **36.2%** | **0.45** | **666.7** | **4.3** | **55** | **24** | **43.05** | **1.79** | **0.42** | **This work**[f] |

[a] The authors report 7.2 ± 0.2, 15.3 ± 0.2, 16.9 ± 0.2 mM release of TPA for Tfu_0883, Tfu_0883(I218A) and Tfu_0883(Q132A/T101A) respectively, after 2 h, corresponding to 1.20, 2.54 and 2.81 g/L of TPA. Regardless of this being in the range of 34-80% of theoretical yield of TPA from PET, the authors do not mention such significant weight loss in the textile material, but rather describe the process as surface modification. [b] Crystallinity determined after melt-extrusion amorphization of post-consumer PET plastic. [c] Determined after 3 rounds of RAging (1 round = 5 min milling at 30 Hz, aging for 24 h at 55˚C). [d] Enzyme is added each round to the remaining PET (0.65 wt%), the concentration reported here in mg $ml^{-1}$ is expressed cumulatively. [e] Determined after 21 cycles of RAging (1 cycle = 5 min milling at 30 Hz, aging for 24 h at 55˚C), washing out TPA and adding 0.65% w/w enzyme every 3 cycles (see Fig. 5, main text). [f] Determined after 1 round of RAging (1 round = 5 min milling at 30 Hz, aging for 24 h at 55˚C).
